# Novel RNA-binding protein YebC enhances translation of proline-rich amino acid stretches in bacteria

**DOI:** 10.1101/2024.08.26.607280

**Authors:** Dmitriy Ignatov, Vivekanandan Shanmuganathan, Rina Ahmed-Begrich, Kathirvel Alagesan, Christian Karl Frese, Chu Wang, Kathrin Krause, Emmanuelle Charpentier

## Abstract

The ribosome employs a set of highly conserved translation factors to efficiently synthesise proteins. Some translation factors interact with the ribosome in a transient manner and are thus challenging to identify. However, proteins involved in translation can be specifically identified by their interaction with ribosomal RNAs. Using a combination of proteomics approaches, we identified novel RNA binding proteins in the pathogenic bacterium *Streptococcus pyogenes*. One of these, a universally conserved protein YebC, was shown to transiently interact with 23S rRNA near the peptidyl-transferase centre. Deletion of *yebC* moderately affected the physiology and virulence of *S. pyogenes*. We performed ribosome profiling and detected increased pausing at proline-rich amino acid stretches in the absence of functional YebC. Further results obtained with *in vivo* reporters and *in vitro* translation system suggest that YebC is a novel translation factor required for efficient translation of proteins with proline-rich motifs.

## Introduction

Bacteria use RNA binding proteins (RBPs) to express and regulate genes^1^. Despite decades of studies on bacterial genetics and physiology, there are still many proteins whose functions remain unknown. Some of these proteins might perform their function by interacting with RNA. Recently, several methods have been developed that enable the identification of bacterial RBPs on a whole-proteome scale. The Grad-seq approach identifies RBPs that form stable complexes with RNA by the transcriptomic and proteomic analysis of fractionated cellular lysates^2^. More transient protein-RNA interactions can be captured by UV crosslinking. The resulting covalently bound protein-RNA complexes can be specifically enriched by purification on silica beads^3^, organic extraction^4, 5^, or pull-down of artificially polyadenylated bacterial transcripts^6^. Subsequent proteomic analysis of the proteins cross-linked with RNA revealed a repertoire of RBPs in *Escherichia coli*^3, 4, 6, 7^, *Salmonella* Typhimurium^5^ and *Staphylococcus aureus*^8^. These studies identified a novel regulatory protein, ProQ^2^, and showed that the glycolytic enzyme enolase in *E. coli* and the transcription factor CcpA in *S. aureus* function as RBPs^3, 8^. A number of proteins of unknown function have also been shown to interact with RNA^5, 6, 7^ and further characterization of these proteins might uncover new aspects of prokaryotic RNA biology.

*Streptococcus pyogenes* is a gram-positive bacterial pathogen causing not only benign human infections such as pharyngitis and impetigo, but also potentially life-threatening invasive diseases such as septicaemia, streptococcal toxic shock syndrome and necrotizing fasciitis^9^. *S.* pyogenes has served as a model organism for RNA biology: the transcriptome analysis has identified multiple non-coding RNAs^10^, the analysis of RNA cleavage by different ribonucleases has provided insights into their modes of action^11^, and the study of the CRISPR-Cas9 system has resulted in the development of new biotechnological tools^12^. In this study, we systematically characterized the RBPs in *S. pyogenes*. Our approach detected most of the annotated RBPs involved in RNA synthesis and degradation, as well as ribosome assembly and translation. In addition, we identified a group of proteins of unknown function that interact with RNA. One of these proteins, YebC, was selected for further characterization.

YebC is a ubiquitous protein present in bacteria and eukaryotic mitochondria. Our results suggest that YebC transiently interacts with 23S rRNA in the vicinity of the peptidyl transferase centre (PTC). Further analysis shows that YebC enhances translation of proline-rich amino acid stretches. The consecutive proline residues impose an unfavourable peptidyl-tRNA geometry in the translating ribosome. In bacteria, the translation elongation factor P (EF-P) enhances translation of polyproline-containing proteins. In the absence of EF-P, the ribosome stalls at polyproline stretches. The interaction of EF-P with the ribosome promotes a geometry favourable for peptide bond formation and alleviates the translational stalling^13, 14, 15^. Recently, an ABCF ATPase YfmR/Uup has also been shown to promote translation of proline-rich motifs in cooperation with EF-P^16, 17, 18^. Our study shows that YebC enhances translation of polyproline stretches *in vivo* and *in vitro*, in the absence of EF-P and YfmR, and therefore represents a novel translation factor.

## Results

### UV-mediated RNA-protein crosslinking identifies RBPs in *S. pyogenes*

The first step of our study was identification of RBPs in *S. pyogenes*. To discover proteins interacting with RNA, we grew *S. pyogenes* to mid-logarithmic growth phase and irradiated the cells with UV light at 254 nm. The cross-linked protein-RNA complexes were specifically enriched using the orthogonal organic phase separation (OOPS) technique^4^ (Fig. 1A and Supplementary fig. 1A). Using stable isotope labelling by amino acids in cell culture (SILAC), we calculated the abundance of proteins in the UV-irradiated relative to non-irradiated samples. The resulting OOPS enrichment values were used as a measure of interaction with RNA (Supplementary data 1 and Supplementary fig. 1B). We further identified amino acids cross-linking with RNA (RNA-binding sites) in *S. pyogenes* proteins using the RBS-ID approach (Supplementary fig. 1C)^19^. In total, our experiment identified 1440 RNA-binding sites in 254 *S. pyogenes* proteins (Supplementary fig. 1D).

**Figure 1.**
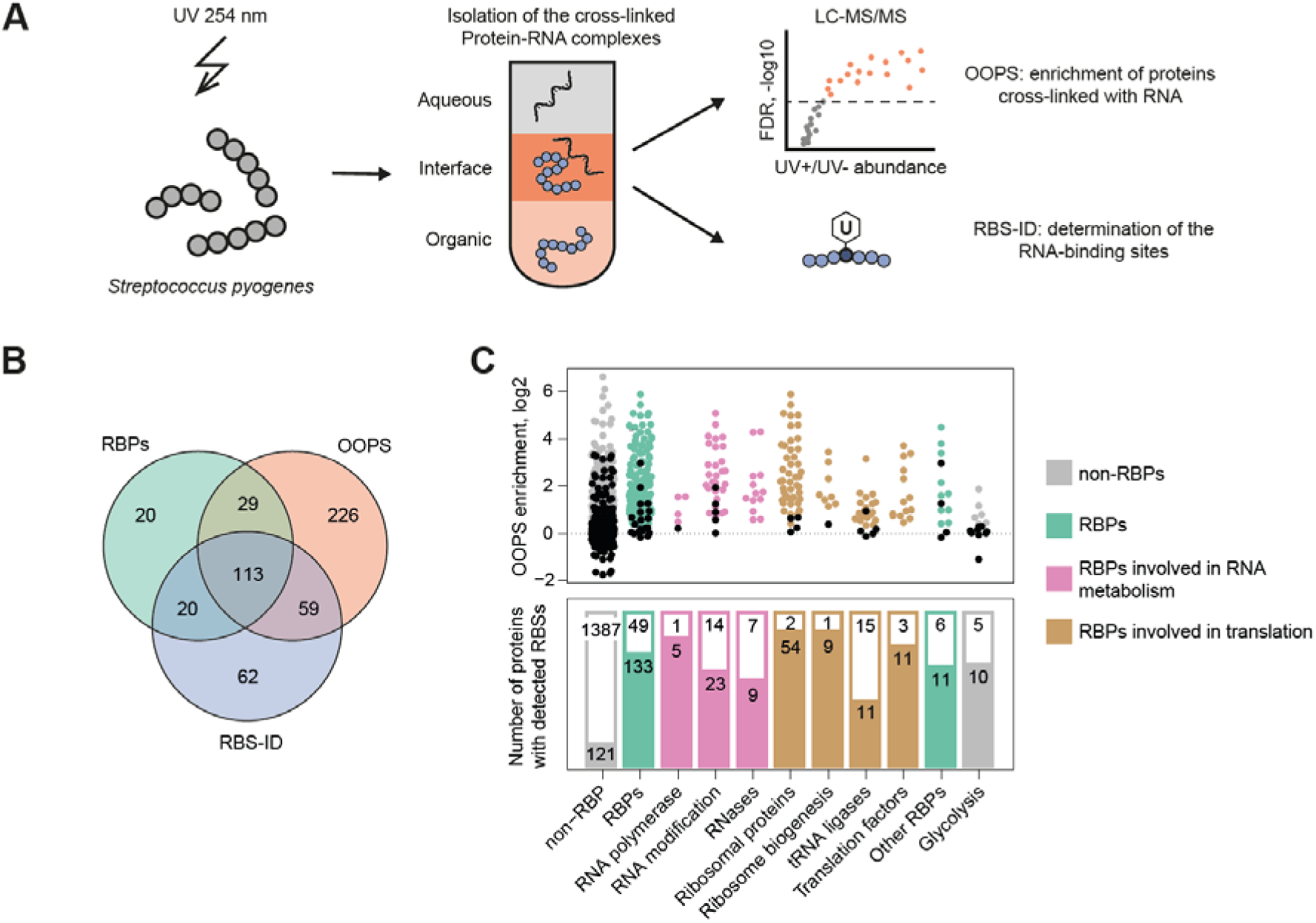
Identification of RBPs in *S. pyogenes.* **(A)** Workflow for the characterization of proteins that cross-link with RNA. After UV 254 nm cross-linking, the protein-RNA complexes were separated from proteins and RNA by acid guanidinium thiocyanate-phenol-chloroform extraction. The cross-linked protein-RNA complexes were further analysed by OOPS or RBS-ID approaches. **(B)** Overlap between annotated RBPs, proteins with statistically significant OOPS enrichment and proteins with detected RNA-binding sites. **(C)** OOPS enrichment and the presence of RNA-binding sites for different functional groups of the annotated RBPs and for glycolysis enzymes. The OOPS enrichment values for proteins in each category are presented in a bee swarm plot. The proteins whose OOPS enrichment is not statistically significant are indicated by black dots. The bar plot shows the number of proteins with identified RNA-binding sites (colored bars) and without RNA-binding sites (white bars) for each category.

Next, we searched the KEGG and Gene Ontology databases to create a list of annotated RBPs for *S. pyogenes*^20, 21^. After manual curation, 182 proteins were annotated as RBPs in *S. pyogenes* (Supplementary data 2). The annotated RBPs, the proteins with significant OOPS enrichment, and the proteins with detected RNA-binding sites overlapped significantly (Fig. 1B). Most of the annotated RBPs in *S. pyogenes* showed statistically significant OOPS enrichment and contain detected RNA-binding sites (Fig. 1C). These include RBPs involved in both RNA metabolism and translation. Glycolysis enzymes from different species often function as non-canonical RBPs^22^. Most proteins in this functional category were not strongly enriched in our OOPS dataset. However, 6-phosphofructokinase, fructose-bisphosphate aldolase, triosephosphate isomerase, two paralogs of phosphoglycerate mutase and L-lactate dehydrogenase were enriched by OOPS, contain RNA-binding sites, and could therefore interact with RNA (Supplementary fig. 2).

### Proteins of unknown function interact with RNA

We hypothesized that some of the OOPS-enriched proteins not annotated as RBPs represent novel RBPs. 59 proteins not annotated as RBPs had statistically significant OOPS enrichment and contained RNA-binding sites (Fig. 1B). We narrowed this selection to 30 proteins using an OOPS enrichment greater than two as an additional criterium (Supplementary fig. 3), and considered these proteins as candidates for novel RBPs (Table 1). Nine of these potential RBPs are annotated as DNA-binding proteins (Fig. 2A). One of them, the transcription factor CcpA, a master regulator of carbon metabolism, has recently been shown to function as a non-canonical RBP in *Staphylococcus aureus*^8^. The other candidates for novel RBPs are several metabolic enzymes and proteins involved in transport that could moonlight as RBPs. One signalling protein, the small alarmone synthase RelQ, has previously been shown to interact with RNA *in vitro*^23^. The protein metabolism factors, peptide deformylase and trigger factor, have been shown to interact with 23S rRNA^24, 25^.

**Figure 2.**
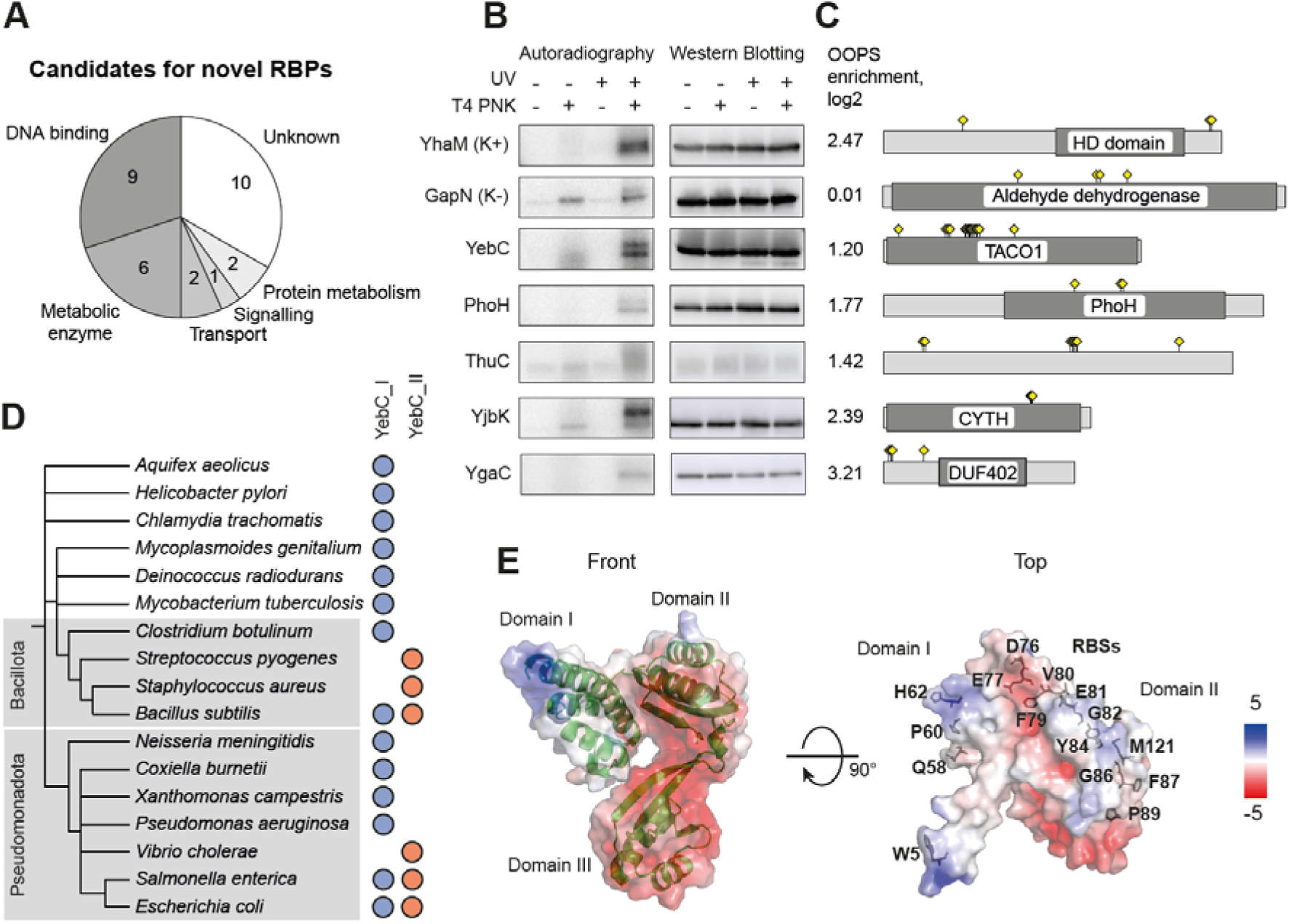
Novel RBPs in *S. pyogenes.* **(A)** Functional categories of candidates for novel RBPs. **(B)** Detection of cross-linked protein-RNA complexes. The strains containing the 3x FLAG tagged potential RBPs or the control proteins YhaM and GapN were UV irradiated. The protein-RNA complexes were immunoprecipitated, radioactively labelled with T4 PNK, separated by gel electrophoresis and transferred to a membrane. Radioactive signals were detected by phosphor imaging. Western blotting with an anti-FLAG antibody served as a control for successful immunoprecipitation. **(C)** Description of proteins in panel B. OOPS enrichment, annotated domains and location of RNA-binding sites are shown. **(D)** Presence of YebC_I and YebC_II variants in different bacteria. **(E)** Modelled structure of YebC in *S. pyogenes*. The cartoon structure of the protein and the predicted surface electrostatic potential are shown. The identified RNA-binding sites are indicated in the top view of the protein.

**Table 1.** Candidates for novel RBPs in *S. pyogenes*. For each RBP candidate, a locus tag, a gene name (if available), a UniProt ID and a manually curated functional annotation are provided along with OOPS enrichment values and identified RNA-binding sites. The asterisks indicate the proteins whose interaction with RNA was confirmed using immunoprecipitation and radioactive labelling of co-precipitated RNA.

| Locus tag | Gene | UniProt ID | Functional category | OOPS enrichment, log2 | RNA-binding sites |
| --- | --- | --- | --- | --- | --- |
| *SPy_0190* |  | Q9A1M3 | DNA binding | 2.182 | 2S; 134W |
| *SPy_0316\** | *yebC* | P67188 | Unknown | 1.199 | 5W; 58Q; 60P; 62H; 76D; 77E; 79F; 80V; 81E; 82G; 84Y; 86G; 87F; 89P; 121M |
| *SPy_0471\** | *phoH* | Q9A145 | Unknown | 1.772 | 177F; 219P; 221A |
| *SPy_0514* | *ccpA* | Q9A118 | DNA binding | 1.307 | 182C |
| *SPy_0534* | *aroE.2* | Q9A102 | Metabolic enzyme | 1.369 | 59S; 61C; 62L; 64D; 65K; 68H; 75C; 109Y; 126Y; 132F; 134I; 135S; 136H; 244P; 246H; 288F |
| *SPy_0539\** | *thuC* | Q9A0Z7 | Unknown | 1.415 | 37Y; 39S; 173L; 175D; 176N; 178L; 179F; 273T |
| *SPy_0598* |  | Q9A0V5 | Metabolic enzyme | 1.868 | 127Y |
| *SPy_0714* | *adcA* | Q9A0L9 | Transport | 1.673 | 59M; 182F; 279E; 289P; 299K; 333T |
| *SPy_0752* |  | Q9A0J4 | Metabolic enzyme | 1.514 | 270C |
| *SPy_0903* | *tcyJ* | Q9A074 | Transport | 1.323 | 116F |
| *SPy_0939* |  | Q9A042 | DNA binding | 1.098 | 221C |
| *SPy_1121* |  | Q99ZR2 | Unknown | 2.145 | 76Q; 77S; 78G; 80R; 81L; 97F; 99F |
| *SPy_1124\** | *yjbK* | Q99ZQ9 | Unknown | 2.386 | 136Y; 137N; 138L |
| *SPy_1125* | *relQ* | Q99ZQ8 | Signalling | 1.529 | 38P; 43T; 44G |
| *SPy_1126* | *nadK* | P65781 | Metabolic enzyme | 3.689 | 21Y; 29K; 30L; 31F; 33V; 38P; 40F; 41Y; 123G; 223R |
| *SPy_1193* |  | Q99ZK4 | Unknown | 1.640 | 15L; 113L; 114Y |
| *SPy_1259* |  | Q99ZE8 | DNA binding | 1.607 | 48S |
| *SPy_1429* | *gpmA* | Q99Z29 | Metabolic enzyme | 1.168 | 113W |
| *SPy_1534* |  | Q99YU4 | Unknown | 2.608 | 206H; 207F |
| *SPy_1582* |  | Q99YR0 | Unknown | 2.835 | 243C |
| *SPy_1607* | *recX* | Q99YP2 | DNA binding | 1.999 | 64Y; 65F; 66L; 67S; 68F; 239A; 252C |
| *SPy_1608\** | *ygaC* | Q99YP1 | Unknown | 3.211 | 6E; 8D; 9F; 38L |
| *SPy_1744* | *accD* | Q99YE0 | DNA binding | 1.764 | 21G; 23V; 24S |
| *SPy_1746* | *fabZ* | P64110 | DNA binding | 1.220 | 137C |
| *SPy_1751* | *fabK* | Q99YD4 | Metabolic enzyme | 1.537 | 212A; 213A; 214K; 215D; 216I; 226G |
| *SPy_1862* | *ybaB* | P67268 | DNA binding | 2.267 | 93F; 98P |
| *SPy_1896* | *tig* | P0C0E1 | Protein metabolism | 1.359 | 45R; 425S |
| *SPy_1936* | *degV* | P67374 | Unknown | 1.082 | 56A; 59H; 164L; 165V |
| *SPy_1958* | *def* | P68771 | Protein metabolism | 1.284 | 36L; 39C; 96E; 99P; 130G; 131C; 147R |
| *SPy_1960* |  | Q99XY5 | DNA binding | 1.884 | 131C |

Ten candidates for novel RBPs are of unknown function. We selected five of them and checked their cross-linking with RNA using immunoprecipitation and radioactive labelling of co-precipitated RNA with T4 polynucleotide kinase (PNK) (Fig. 2B and 2C)^26^. To this end, we introduced the 3x FLAG tags to the C-termini of their genes. Expression from the native promotors enabled us to estimate the interaction with RNA at physiological concentrations of the proteins. The positive control gene *yhaM* and the negative control gene *gapN* were also tagged with the 3x FLAG tag. The exoribonuclease YhaM has been shown to form stable complexes with RNA in *S. pneumoniae*^27^. Consistent with this, we observed the radioactive signal originating from the co-immunoprecipitated RNA only upon UV cross-linking and in the presence of PNK. For the negative control protein GapN, we also observed some signal in the UV+ PNK+ sample, but it was only marginally higher than in the UV-PNK+ sample. Therefore, a strong increase in radioactive signal after UV cross-linking and in the presence of PNK indicates that a protein interacts with RNA. The proteins YebC and YjbK were expressed at relatively high levels. In the UV-irradiated and PNK-labelled samples, they showed radioactive signals comparable to those of a positive control. These results suggest that YebC and YjbK are *bona fide* RBPs. The proteins YgaC and ThuC were expressed at low levels under the conditions tested. However, specific radioactive signals from cross-linked RNA were also observed for these proteins. Another candidate, the protein PhoH, showed weak cross-linking with RNA. These results validate our approach to the search for novel RBPs and show that most of them interact with RNA.

YebC is a ubiquitous protein found in most bacterial species, suggesting that it plays an important role in bacterial physiology. Nevertheless, the function of YebC remains enigmatic. Our results showed that this protein interacts with RNA, and studying this protein could uncover a novel molecular biology mechanism. Consequently, we focused on YebC with the aim of elucidating its function. The mitochondrial homolog of y*ebC* - a translational activator of cytochrome oxidase subunit I (*TACO1*) - is a nuclear-encoded gene whose product is targeted to the mitochondria. Mutations in *TACOI* in humans and mice result in defective translation of a mitochondrially encoded COXI protein, suggesting that the protein acts as a translational activator^28, 29^.

In bacteria, it has been proposed to divide YebC homologs into two variants: YebC_I and YebC_II^30^. The genomes of some bacteria, such as *E. coli* and *B. subtilis*, encode both variants, while *S. pyogenes* possesses only the YebC_II variant (Fig. 2D). The sequence and structure of YebC are evolutionarily conserved (Supplementary fig. 4). We modelled the structure of YebC from *S. pyogenes* on the basis of YebC from *Coxiella burnetii*^31^. The protein consists of three domains: domains I and II contain the positively charged surface patches, while domain III is highly negatively charged. Interestingly, the RNA-binding sites identified in our experiment are located near the positively charged surfaces, suggesting that they form the interface for interaction with RNA (Fig. 2E).

### Deletion and site-directed mutagenesis of YebC in *S. pyogenes*

We constructed a strain of *S. pyogenes* with a deleted *yebC* gene and a complemented strain in which *yebC* with a C-terminal 3x FLAG tag was reintroduced on an integrative vector under the control of its native promoter. The deletion of *yebC* slightly affected *S. pyogenes* growth: while no difference could be observed in THY or C medium, the Δ*yebC* strain grew to a higher optical density in chemically defined medium (CDM) (Supplementary fig. 5A). Transcriptome analysis performed in THY medium revealed a moderate effect of the *yebC* deletion on gene expression (Supplementary data 3, Supplementary fig. 5B and 5C). Notably, *yebC* deletion significantly decreased expression of the major *S. pyogenes* virulence factor, the secreted protease SpeB^9^ (Fig. 3A), and led to reduced intracellular survival in human macrophages (Fig. 3B and Supplementary fig. 6). In addition, while no difference in *S. pyogenes*-induced cytotoxicity could be observed between the *WT*, Δ*yebC* or Δ*yebC / yebC+* strains (Supplementary fig. 6A), there was a mild albeit not significant reduction of IL-1β and IL-18 secretion by macrophages infected with the Δ*yebC* strain (Supplementary fig. 6B). In contrast, levels of IL-6 remained unaffected.

**Figure 3.**
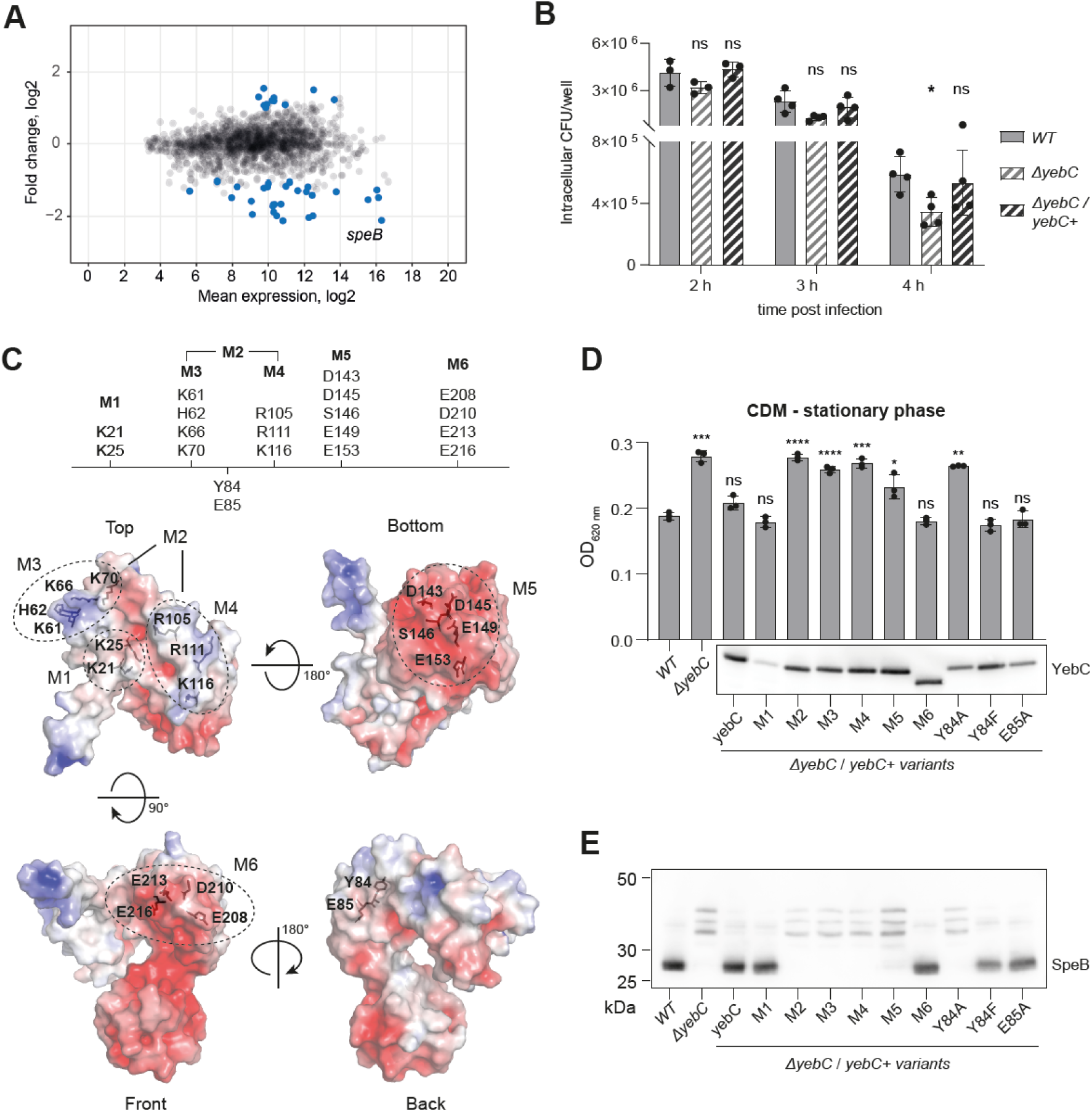
Mutagenesis of YebC in *S. pyogenes*. **(A)** MA plot for the difference in gene expression between Δ*yebC* and *WT* strains in Stat growth phase in THY. Gene expression in the Δ*yebC* strain was compared to *WT* and Δ*yebC* / *yebC+* strains. Genes showing statistically significant changes in both comparisons are marked in blue. *speB* was among the genes with significantly decreased expression in the Δ*yebC* strain. **(B)** Intracellular survival of *S. pyogenes WT*, Δ*yebC*, and Δ*yebC* / *yebC+* in primary human macrophages at 2h, 3h, and 4h post-infection. Data represent the mean ± SD of three biological replicates. Statistical analysis was performed using two-way ANOVA (*p ≤ 0.05). **(C)** Amino acid residues for site-directed mutagenesis superimposed on the YebC structure. **(D)** Optical density of stationary phase cultures of *yebC* mutant and complemented strains in CDM. Data represent mean ± SD of three biological replicates. The optical density of cultures of each strain was compared to that of the *WT*. The statistical significance of the differences was estimated using the *t*-test and the results are indicated above the bars (****p ≤ 0.0001, ***p ≤ 0.001, **p ≤ 0.01, *p ≤ 0.05). The complemented YebC is tagged with the C-terminal 3x FLAG tag and its expression was measured by western blotting using anti-FLAG antibodies (shown below). **(E)** Expression of the SpeB protein in the *yebC* mutant and complemented strains. The strains were grown overnight in THY, the supernatants were collected, and SpeB expression was probed with anti-SpeB antibodies. In the culture supernatant of the *WT* strain, SpeB was present as a mature 28 kDa enzyme, while in the supernatant of *yebC* mutant, the bands corresponding to the 40-kDa zymogen and several intermediates were detected.

Next, we used site-directed mutagenesis to identify amino acids important for YebC function. Several groups of conserved amino acids forming positive and negative patches on the protein surface were selected (Fig. 3C) and the amino acids were substituted with alanine. The mutated versions of *yebC* were introduced into the Δ*yebC* strain. The deletion of *yebC* in *S. pyogenes* was manifested by an increase in the optical density of the stationary phase culture in CDM (Fig. 3D). Consistent with the transcriptomics data, the level of SpeB in the culture supernatant of the Δ*yebC* strain also decreased (Fig. 3E)^32^. Complementation with the wild type y*ebC* restored both phenotypes. Some of the mutated versions of *yebC* were unable to restore the phenotypes and these mutations were considered to inactivate the protein.

The M1 mutation is a substitution of the highly conserved K21 and K25 residues by alanine residues. This mutation significantly reduced the level of YebC, probably by affecting the protein’s stability. Nevertheless, the M1 mutant protein was able to fully complement the Δ*yebC* phenotype, showing that the M1 mutation does not inactivate the protein. The M2 mutation comprises several conserved amino acids that form two positively charged patches on the surface of YebC. This mutation makes *yebC* unable to restore the phenotype. With the M3 and M4 mutations, we separately substituted the groups of amino acids that form these positively charged patches. The mutant versions of *yebC* were also not able to complements the phenotype, suggesting that the positively charged amino acids located on the upper part of the protein are essential for its function.

The *yebC* gene with the mutation M5 also did not complement the phenotype, suggesting the importance of the conserved amino acids forming the negatively charged surface at the lower tip of domain III. In contrast, the M6 mutation, which affects several negatively charged amino acids in domain II, had no effect on YebC activity. Amino acids Y84 and E85 are universally conserved and Y84 represents an RNA-binding site (Fig. 2E). The *yebC* gene with tyrosine 84 substituted by alanine, but not by phenylalanine, did not restore the phenotype, suggesting that the tyrosine’s aromatic ring, but not its hydroxyl group, is crucial for YebC activity. Substitution of glutamic acid 85 by alanine had no effect on YebC activity and this amino acid residue appears to be dispensable. In conclusion, the site-directed mutagenesis identified the importance of the positively charged amino acids at the top of domains I and II, and of the universally conserved Y84 for YebC activity.

### YebC transiently interacts with the ribosome

We next used the iCLIP approach to study the interaction of YebC with RNA. The YebC variant with the C-terminal 3x FLAG tag was able to fully complement the Δ*yebC* phenotype (Fig. 3D), suggesting that the tag does not interfere with YebC function. We irradiated the *yebC:3xFLAG* strain with UV 254 nm light. The cross-linked YebC-RNA complexes were immunoprecipitated, the RNA was partially degraded with RNase I (Supplementary fig. 7A) and cDNA libraries were prepared according to the iCLIP protocol^33^. In parallel, the control libraries were prepared from UV-irradiated *WT* and non-irradiated *yebC:3xFLAG* strains. As a control for non-specific background, we also prepared size-matched input (SMI) libraries^34^. To analyse the iCLIP data, we extracted the cross-linked nucleotides located immediately upstream of the mapped cDNA reads^35^ and mapped the regions with increased number of the cross-linked nucleotides (Supplementary data 4).

More than 20% of the cross-linked nucleotides in the *yebC:3xFLAG* UV+ sample were located in a single region of 23S rRNA (Fig. 4A). No other region in this sample or in any of the controls contained such a high proportion of the cross-linked nucleotides. This locus of 23S rRNA was enriched for the cross-linked nucleotides only in the *yebC:3xFLAG* UV+ sample and not in the controls (Fig. 4B), suggesting that it represents the binding site for YebC. Visual inspection of the cross-linked nucleotides and regions did not identify any other locus significantly enriched in *yebC:3xFLAG* UV+ compared with the controls. Therefore, YebC likely represents a protein that interacts specifically with 23S rRNA. The cDNA fragments mapping to this region encompass helix 89, and most of them terminate directly upstream of nucleotides A2453, A2454 and A2456, which apparently represent the the cross-linked nucleotides (Fig. 4C). These nucleotides are homologous to A2450, A2451 and A2453 in *E. coli*. They are part of the PTC and are essential for the ribosome function^36^.

**Figure 4.**
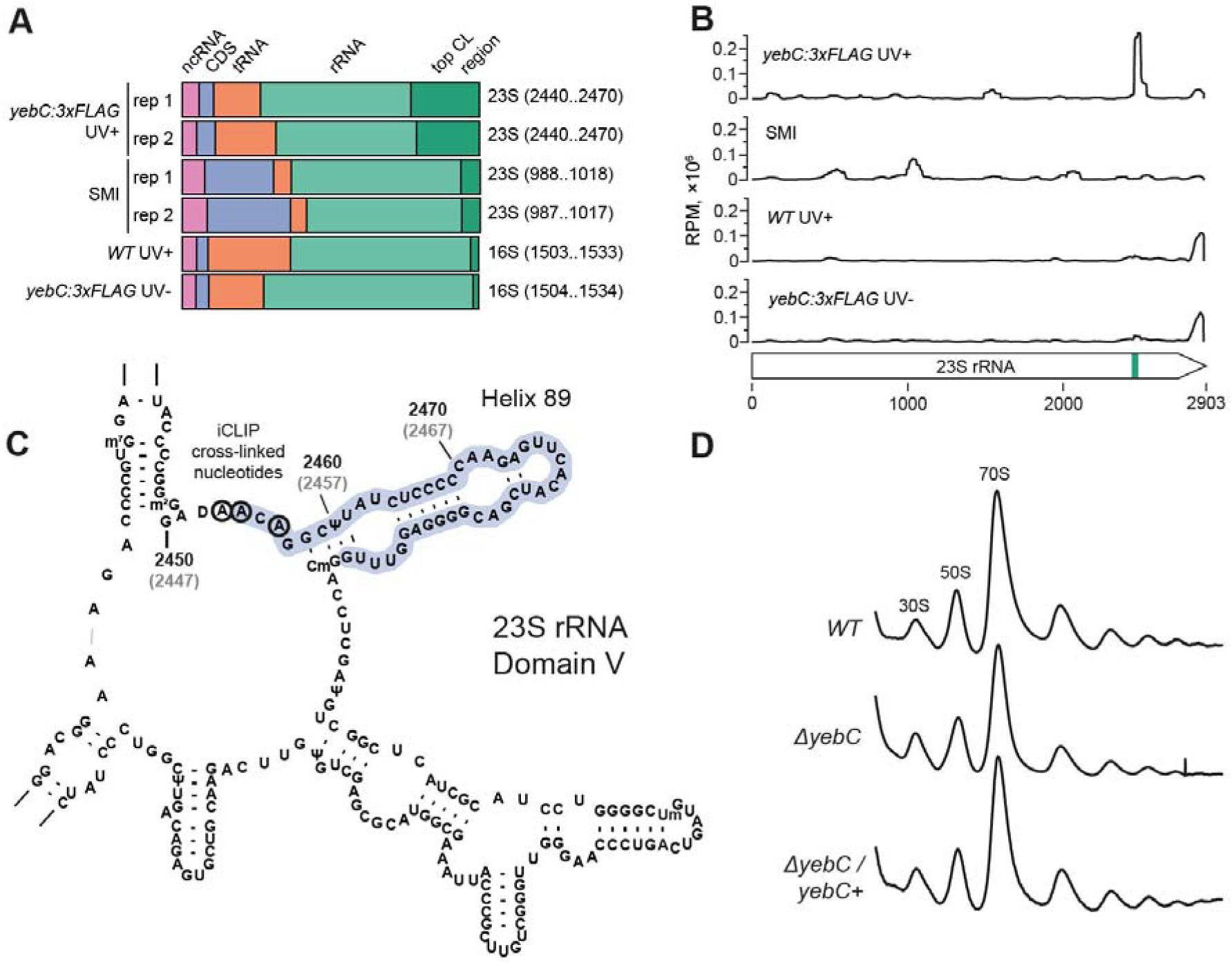
Interaction of YebC with the ribosome. **(A)** Distribution of the nucleotides cross-linked to YebC in the *S. pyogenes* transcriptome. Genomic regions (30 nt in length) showing an increased number of cross-linked nucleotides relative to neighboring regions were determined. The region with the highest number of cross-linked nucleotides for each library is highlighted on the right. **(B)** iCLIP read coverage of 23S rRNA in *yebC:3xFLAG* UV+ and control samples. Coverage was normalised to the sequencing depth of the libraries and presented as RPM values. The top cross-linking region in *yebC:3xFLAG* UV+ encompassing nucleotides 2440 to 2470 is marked in green on the 23S rRNA. **(C)** Position of the cross-linking region in domain V of 23S rRNA. The nucleotides with increased iCLIP read coverage are coloured in blue and the cross-linked nucleotides are marked with black rings. The coordinates of nucleotides in the *S. pyogenes* genome are shown in black and the coordinates of the homologous nucleotides in *E. coli* are shown in grey. **(D)** Sucrose density gradient ribosome traces of *WT*, Δ*yebC* and Δ*yebC / yebC+* strains.

Our search for RBPs in *S. pyogenes* identified most of the proteins involved in translation, including ribosomal proteins, proteins involved in ribosome biogenesis and translation factors. Therefore, it seemed possible that YebC is also involved in ribosome biogenesis or translation. In order to investigate the contribution of YebC to ribosome biogenesis, we resolved ribosomes isolated from mid-logarithmic and stationary growth phase *S. pyogenes* cultures on sucrose density gradients and demonstrated that YebC does not form a stable complex with translating ribosomes or free ribosomal subunits (Supplementary fig. 7B). Deletion of *yebC* did not affect the polysome profile (Fig. 4D), suggesting that the protein is not involved in ribosome maturation. In conclusion, our data indicate that YebC transiently interacts with the ribosome at or near the PTC and Helix 89 and therefore may affect the peptide bond formation.

### In the absence of YebC, the ribosome pauses at proline-rich regions

To study how the mutation of *yebC* affects translation, we used the ribosome profiling approach. The strains with functional YebC (*WT* and Δ*yebC / yebC+*) and with YebC mutation (Δ*yebC* and Δ*yebC / yebC_M2+*) were grown to the mid-logarithmic growth phase and collected by rapid filtration to preserve the native positions of the translating ribosomes^37^. To accurately identify the ribosome positions, we selected the 27-40 nucleotide long ribosome-protected mRNA fragments and mapped the ribosomal P sites 15 nucleotides upstream of the 3′ ends of these mRNA fragments (Supplementary fig. 8A and 8B)^38^. For each codon, we calculated the pause score, defined as the number of reads mapped to a particular position divided by the average read density for the corresponding gene. Median pause scores were slightly higher when serine or glycine codons were located in the E site (Supplementary fig. 8C). This amino acid specific increase in pausing is explained by the use of chloramphenicol to stop translation in the lysate^39^ and does not depend on the presence of functional YebC.

The ribosome pause scores of the Δ*yebC* and Δ*yebC / yebC_M2+* strains were similar to each other but different from the strains with the functional YebC, further demonstrating that the M2 mutation inactivates YebC (Supplementary fig. 9). We searched for codons where the ribosome pausing changes upon YebC mutation. In total, we identified 168 positions with a statistically significant difference in pause score (Supplementary data 5). For 153 of these positions, the ribosome pausing was increased in the strains with inactive YebC. Interestingly, pausing was strongly increased in the vicinity of a triple proline codon in the *valS* mRNA (Fig. 5A). Analysis of the genomic context demonstrated that the ribosome pausing was often increased directly at and downstream of proline-rich amino acid sequences (Fig. 5B). The codons located in close proximity to each other were often affected, and we identified a total of 81 positions across the transcriptome showing increased pausing in *yebC* mutants. Most of the loci were located in the vicinity of the PP, PXP or DXP amino acid motifs. Among these, the sequences PPG, PIP and DIP were overrepresented (Fig. 5C and Supplementary data 6). We searched for these amino acid sequences in *S. pyogenes* proteins and calculated the average pause score in their vicinity (Fig. 5D). In the Δ*yebC* strain, the pausing score at PPG increases when the second proline is located in the P site and remains elevated for three codons after PPG translation. The increase in pausing score for PIP and DIP is not as pronounced, suggesting that these motifs induce a weaker ribosome pausing. However, in this case, pausing also continues even when the ribosome translates several codons after the motifs.

**Figure 5.**
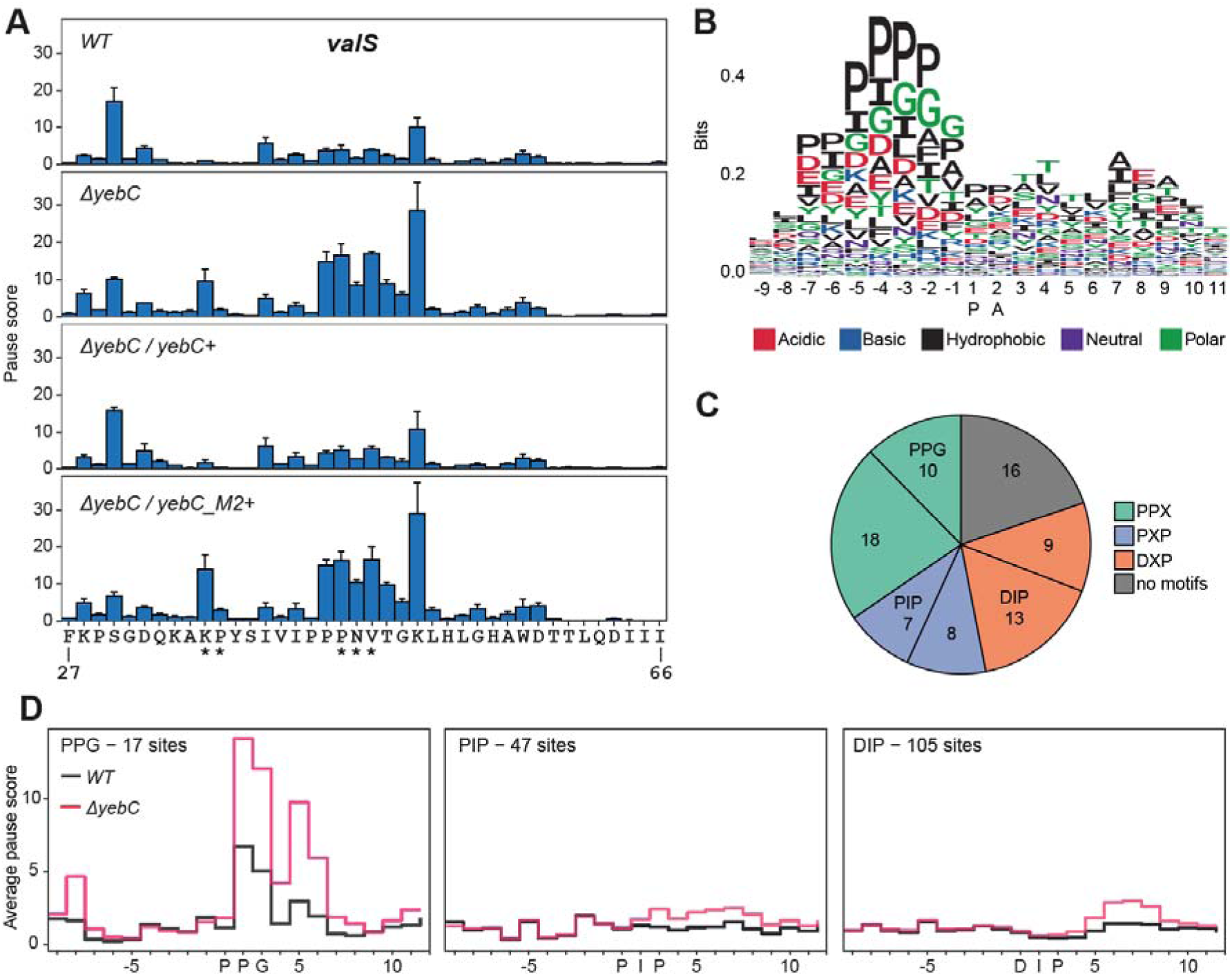
Ribosome pausing in *S. pyogenes yebC* mutants. **(A)** Mean pause scores for *valS* codons 27-66 in *WT* and *yebC* mutant strains. Data represent mean ± SD of three biological replicates. Codons with a statistically significant difference in pause scores between the strains with functional and non-functional YebC are indicated by asterisks. **(B)** Logo plot of codons in the vicinity of the sites with increased pausing in *yebC* mutant strains. **(C)** Motifs enriched in the vicinity of the sites with increased pausing in *yebC* mutant strains. **(D)** Average pausing scores for the codons surrounding PPG, PIP and DIP motifs. The numbers of these motifs in *S. pyogenes* mRNAs are indicated.

### YebC rescues ribosome stalling at proline-rich regions

Our ribosome profiling data suggest that YebC alleviates ribosome stalling during translation of the proline-rich amino acid stretches. We examined how this affects the expression of polypeptides with the polyproline sequences. To this end, we constructed N-terminal 3x FLAG-tagged sfGFP-mKate fusions with different proline-rich sequences in the linker (Fig. 6A) and introduced them to the *WT* and Δ*yebC* strains of *S. pyogenes*. Expression of the reporter proteins was induced for a short period of time and analysed by western blotting with anti-FLAG antibodies (Fig. 6B). In the *WT* strain, the reporter protein expression was not affected by the introduction of the proline-rich sequences. In contrast, expression of the reporter protein with P5 and P3 motifs was significantly downregulated in the Δ*yebC* strain.

**Figure 6.**
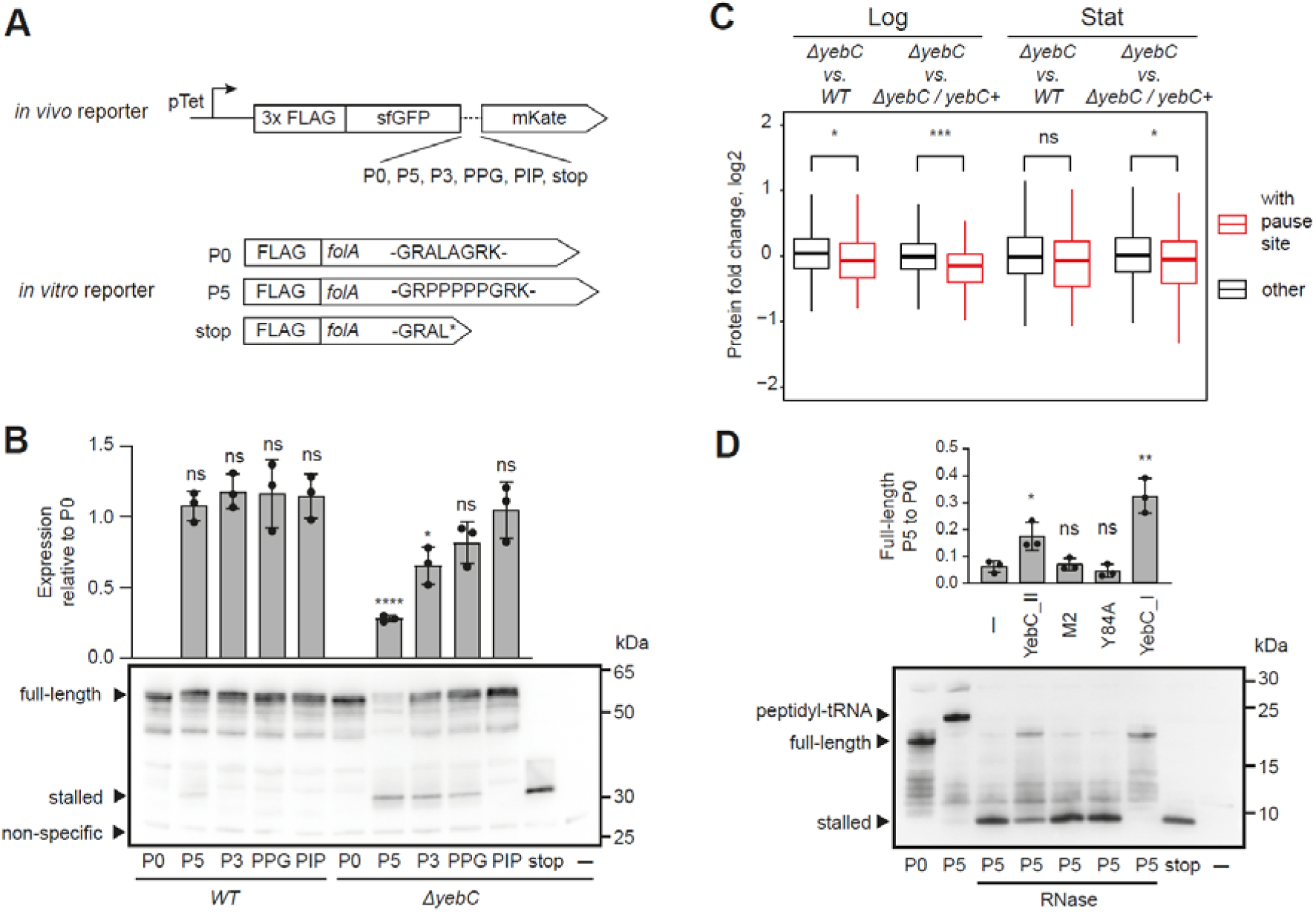
Effect of YebC on translation of proline-rich regions. **(A)** Schematic of *in vivo* and *in vitro* reporters for ribosome stalling. **(B)** Effect of *yebC* deletion on the polyproline stalling *in vivo*. The reporter proteins bearing different amino acid sequences between sfGFP and mKate (panel A) were introduced to the *WT* and Δ*yebC* strains under the control of the Ptet promoter. Expression of the reporter was induced in the exponentially growing *S. pyogenes* cells and the cells were collected 30 minutes after induction. Western blotting with anti-FLAG antibodies detected the full-length reporter, the truncated reporter with stop codon and the truncated reporters in the Δ*yebC* strain. The non-specific band served as a loading control. The amount of the full-length products relative to P0 was measured by densitometry and presented in the barplot as mean ± SD of three biological replicates. The statistical significance of the difference relative to the P0 strain was estimated with a *t*-test (****p ≤ 0.0001, *p ≤ 0.05). **(C)** Genes with YebC-dependent pausing are expressed at a lower level in the Δ*yebC* strain. The difference in protein expression in the Δ*yebC* compared with the *WT* and Δ*yebC / yebC+* strains was measured by MS proteomics. The distribution of the changes for genes with and without YebC-dependent pause sites is presented as a boxplot, where the box represents the interquartile range and the median and the whiskers extend to ×1.5 of the interquartile range. The statistical significance of the difference in protein expression was estimated with the Mann-Whitney test (***p ≤ 0.001, *p ≤ 0.05). **(D)** Effect of YebC on the polyproline stalling *in vitro*. The *folA* mRNA with the indicated mutations served as a template for *in vitro* translation using the PURE system. The *E. coli* YebC proteins b1983 and b1864 and the mutant versions of b1983 were added to the reactions with P5 mRNAs (panel A). The indicated reactions were then treated with RNase A. The barplot represents the amount of the full-length product in P5 measured by densitometry and normalised to P0 as mean ± SD of three replicates. The significance of the effect of YebC proteins on the amount of the full-length product was estimated with the *t*-test (**p ≤ 0.01, *p ≤ 0.05).

The PPG linker also caused a slight downregulation, which was however not statistically significant, and the PIP linker had no effect. Interestingly, in the Δ*yebC* strain, we also observed reporters with P5, P3 and PPG linkers that were truncated at the proline-rich stretches. The truncated products probably appear when the ribosome stalls at these amino acid stretches and is unable to synthesise the full-length proteins.

This experiment suggested that YebC is essential for the efficient translation of proteins containing polyproline stretches. We wanted to examine how the increased ribosome pausing in the absence of YebC affects protein expression at the whole-proteome level. To this end, we performed a proteome analysis of the same bacterial cultures that we used for the transcriptome analysis: *WT*, Δ*yebC* and Δ*yebC / yebC+* strains in mid-logarithmic and stationary growth phases (Supplementary data 7). We compared the protein levels in the Δ*yebC* strain with those in the *WT* and Δ*yebC / yebC+* strains (Fig. 6C). Interestingly, the proteins with YebC-dependent ribosome pause sites demonstrated a slight but statistically significant down-regulation in the Δ*yebC* strain in the logarithmic growth phase. In the stationary growth phase, we also observed a trend towards downregulation, but it was not statistically significant.

We next used the PURE *in vitro* translation system^40^ to test whether YebC is able to rescue the ribosome stalling at the polyproline amino acid stretches *in vitro*. This is a reconstituted system in which all components are purified from *E. coli*. As a result, the PURE system enables the effect of YebC on translation to be monitored in the absence of additional factors. Unlike *S. pyogenes*, the genome of *E. coli* encodes two versions of *yebC* (Fig. 2D). To test the difference between these versions, we expressed and purified *E. coli* YebC_I and YebC_II. We also expressed and purified *E. coli* YebC_II with the M2 and Y84A mutations, which have been shown to inactivate the protein’s activity in *S. pyogenes*. As templates for *in vitro* translation, we used mRNAs encoding mutant versions of the FolA protein with an N-terminal FLAG tag (Fig. 6A). Translation of the P0 version produced a full-length protein of 18.9 kDa (Fig. 6D). In contrast, translation of the P5 version resulted in a product of higher mass. After treatment with RNase A, the mass of the P5 product decreased to approximately 7.1 kDa - the mass of FolA with a premature stop codon in place of the P5 sequence. These data indicate that ribosomes almost completely stall at the pentaproline amino acid stretch and are unable to synthesise full-length FolA. The addition of YebC_II to the reaction partially rescued the ribosome stalling and enabled the full-length protein to be synthesised, albeit with a much lower efficiency.

Importantly, the M2 and Y84A mutants were completely inactive, demonstrating the specificity of YebC_II activity in our system. Interestingly, YebC_I showed higher activity than YebC_II: the stalled product was not visible and the amount of full-length protein was higher. The difference in activity observed between YebC_I and YebC_II suggests that these proteins are functionally divergent. In conclusion, our experiments with *in vivo* and *in vitro* reporters demonstrate the ability of YebC to alleviate ribosome stalling on polyproline amino acid stretches.

## Discussion

In this study, we used the OOPS and RBS-ID approaches to identify RBPs in *S. pyogenes*. A combination of these techniques reliably identified the majority of annotated RBPs. A group of proteins not previously shown to interact with RNA also showed statistically significant OOPS enrichment. OOPS enrichment greater than two and the presence of identified RNA-binding sites were additional criteria for the identification of novel RBPs. This significantly narrowed the set of candidates for novel RBPs and filtered out the proteins that could stably interact with polysaccharides^4^.

Our list of candidates for novel RBPs in *S. pyogenes* includes 30 proteins (Table 1). Analysis of the literature shows that four of these proteins have homologs in other bacteria that interact with RNA^8, 23, 24, 25^, confirming our data. Ten of the candidates for novel RBPs have an unknown function. We selected five of these and tested their cross-linking with RNA using immunoprecipitation and radioactive labelling of co-immunoprecipitated RNA. The protein PhoH showed weak cross-linking to RNA. The PhoH domain belongs to the P-loop NTPases and has been proposed to function as an RNA helicase^41^. The other potential RBPs, YgaC and ThuC, also showed cross-linking to RNA, but were expressed at low levels under the condition tested. YgaC contains the DUF402 domain and its homolog in *S. aureus* possesses nucleoside diphosphatase activity^42^. ThuC is one of the proteins encoded by a biosynthetic gene cluster probably involved in the synthesis of a secondary metabolite^43,44^. Two other potential RBPs, YjbK and YebC, were highly expressed and exhibited robust cross-linking with RNA. YjbK belongs to the CYTH domain superfamily - the metal-dependent phosphohydrolases, whose active site is located within a topologically closed hydrophilic beta-barrel. The representatives of this superfamily have diverse activities. In fungi and protozoa, it is an RNA triphosphatase that catalyses the first step of RNA capping; in mammals, it is a thiamine triphosphatase; and in *E. coli*, a CYTH-domain protein has demonstrated triphosphatase activity^45, 46, 47^.

Among the novel RBPs, we selected YebC for further characterisation. In a recent study, YebC was also shown to interact with RNA in *E. coli*, corroborating our data^6^. Although the function of YebC is not clear, a number of studies suggest that it is involved in translation. The mitochondrial YebC homologs, TACOI in mice and DPC29 in *Saccharomyces cerevisiae*, have been shown to associate with the mitochondrial ribosome. In humans and mice, TACOI is required for efficient translation of the COXI protein, and its absence leads to deficiency of respiratory complex IV^29, 48^. Two studies performed in bacteria also hint at a role of YebC in translation. In *E. coli*, overexpression of YebC rescued the growth defect caused by inactivation of the ribosome-associated GTPase BipA^49^. In *B. subtilis*, overexpression of YebC rescued the defect of flagella assembly caused by deletion of the translation elongation factor P (EF-P). Cells lacking EF-P were defective in hook completion due to translation pausing at a specific PP motif within the flagella protein FliY^50^. While these studies suggest that YebC is involved in translation and is probably necessary for the translation of polyproline motifs, another group of studies suggests that the protein may function as a transcription factor^51, 52, 53, 54, 55^.

YebC is an evolutionarily conserved protein with very similar structures in species as evolutionarily distantly related as the thermophilic bacterium *Aquifex aeolicus* and the mouse^29^. In this study, we show that amino acids that form positively charged surfaces on the top of domains I and II are essential for YebC activity. These surfaces are located in close proximity to the identified RNA-binding sites and could therefore be involved in the interaction with RNA. We also tested the invariant Y84 and E85 positions. Surprisingly, mutation of E85 did not affect the ability of the protein to complement the Δ*yebC* phenotype. For Y84, substitution by an alanine, but not a phenylalanine, was deleterious to YebC activity in the *in vitro* translation system. Given that Y84 represents an RNA-binding site, we hypothesize that the aromatic ring of the tyrosine residue is important for the interaction with RNA.

Our data suggest that YebC interacts with the ribosome: according to iCLIP results, the protein cross-links with the domain V of 23S rRNA. iCLIP relies on reverse transcriptase stopping at the cross-link site between the RNA and peptides remaining of the digested protein. Consequently, YebC is likely to cross-link with the nucleotides constituting the PTC: A2453, A2454 and A2456. However, in some cases reverse transcriptase does not terminate synthesis at the cross-linked nucleotides^56^, and we cannot exclude the possibility that YebC cross-links with a nucleotide located in helix 89 of 23S rRNA. Despite the robust cross-linking, YebC does not appear to form a stable complex with the ribosome and therefore their interaction is rather transient. Some translation factors also interact with the ribosome in a transient manner, and isolation of the interaction intermediate may require the mutant version of a protein or the addition of antibiotics^57, 58^.

The discovery of the interaction with the ribosome prompted us to examine the effect of *yebC* deletion on translation. Ribosome profiling revealed increased pausing at the proline-rich amino acid stretches in the *yebC* mutant. This is similar to the pausing induced by *efp* deletion^38, 50^. However, in the case of YebC, pausing was often induced by the PIP and DIP motifs, which is not typical for *efp* mutants. In addition, the ribosome often pauses a few codons after translation of the polyproline motif in the *yebC* mutant. This suggests that YebC and EF-P have different mechanisms of action. Experiments with *in vivo* reporters in *S. pyogenes* have shown that rescue of ribosome stalling by YebC is necessary for efficient translation. The stalling was induced by the consecutive prolines but not by the PIP motif. According to the ribosome profiling data, only a subset of PIP motifs induce pausing, and this pausing is generally weaker than that at PPG motifs. Therefore, detectable stalling at the PIP motif may only occur in a specific context.

The genes with increased ribosome pausing in the Δ*yebC* strain tend to be expressed at slightly lower levels. In general, the deletion of *yebC* had a pleiotropic effect on the physiology of *S. pyogenes*. The most obvious phenotypes were slightly altered growth and reduced expression of the virulence factor SpeB. Expression of SpeB is regulated by a quorum sensing system and is additionally modulated by environmental pH^59, 60^. We did not identify a YebC-dependent pausing in *speB* or its regulator, and the observed deficiency in SpeB expression is probably an indirect effect of *yebC* deletion. Consistent with SpeB playing a role in intracellular persistence^61^ and cleavage of pro-IL-1β^62^, the strain with deleted *yebC* showed reduced survival in human macrophages and elicited a slight decrease in the release of IL-1β and IL-18.

Our *in vivo* data showed that the effect of *yebC* deletion on translation is similar to the effect of *efp* deletion^38, 50^. To exclude the possibility that YebC function is dependent on EF-P or YfmR, we used the PURE *in vitro* translation system. This experiment showed that both YebC paralogs from *E. coli* are capable of resolving ribosomal stalls on a polyproline stretch. Importantly, the PURE system contains neither EF-P or YfmR, and our results demonstrate the activity of YebC in the absence of these factors. Our data also highlight the difference of activity between YebC_I and YebC_II and suggest that a certain degree of functional diversification exists between YebC variants.

During the preparation of the manuscript, a study demonstrating the role of TACOI in promoting translation of the polyproline motifs in mitochondria was published^63^. Consequently, YebC and its mitochondrial homolog TACOI appear to perform the same function. It is possible that the promotion of translation of proline-rich motifs is not the only function of YebC and this protein has other activities. The molecular mechanism of action for YebC is not clear and its elucidation may require structural studies. Our data suggest that in bacteria, YebC interacts with the ribosome in a transient manner and the isolation of their complex for structural studies may therefore be a non-trivial task. However, it is likely that YebC interacts with the PTC: in *S. pyogenes*, the protein cross-links with the PTC nucleotides, and in mitochondria, it has been proposed that TACOI stabilises the PTC during translation together with the ribosomal protein bL27m^63^. It is remarkable that in bacteria, the ribosome requires the action of three specialised proteins - EF-P, YfmR and YebC - for efficient translation of proline-rich motifs. Future studies should address how these proteins act in a coordinated manner.

## Methods

### *S. pyogenes* cell culture

*Streptococcus pyogenes* strain SF370; M1 GAS (ATCC® 700294™) and its derivatives were cultured on Trypticase soy agar (TSA, BD Difco) plates supplemented with 5% defibrinated sheep blood. In liquid media, the bacteria were cultured statically at 37°C and 5% CO_2_. THY medium (Bacto Todd Hewitt Broth (Becton Dickinson) complemented with 0.2% yeast extract), C medium (0.5% w/v Protease Peptone No. 3 (Difco), 1.5% (w/v) yeast extract (Servabacter) and 1 g/L NaCl) and the chemically defined medium (CDM) were used as liquid media^64^. CDM without L-arginine and L-lysine was prepared from powder (Alpha biosciences # C03-217) and supplemented with 1 mg/L Fe(NO_3_)_3_ · 9 H_2_O, 5 mg/L FeSO_4_ · 7 H_2_O, 3.4 mg/L MnSO_4_ · H_2_O, and 1% glucose. For SILAC labelling, the medium was supplemented with 0.125 mg/mL ^13^C^15^N-labelled L-Lysine HCl (Silantes) and ^13^C^15^N-labelled L-Arginine HCl (Silantes). Alternatively, the non-labelled L-Lysine and L-Arginine HCl were added. Prior to inoculation, NaHCO_3_ and L-cysteine were added to final concentrations of 2.5 and 0.71 g/L, respectively.

### Mutagenesis of *S. pyogenes*

The introduction of 3x FLAG to the C-termini of *S. pyogenes* genes was performed using the Cre-Lox recombination system^65^. For each gene, the sequences upstream and downstream of their stop codons were amplified from *S. pyogenes* genomic DNA with primers listed in Supplementary table 1 and cloned into the suicidal plasmid pSEVA141_3xFLAG-lox71-*erm*-lox66^66^ (Supplementary table 2 and Supplementary fig. 10). The plasmids were introduced by electroporation into *S. pyogenes* and clones with the introduced lox71-*erm*-lox66 cassette were selected. For removal of the cassette, the clones were transformed with the replicative plasmid encoding Cre recombinase^67^. The deletion of the *yebC* gene was also performed with the Cre-Lox recombination system. The sequences upstream and downstream of *yebC* (*SPy_0316*) were cloned into the suicidal plasmid pSEVA141_lox71-*erm*-lox66, the plasmid was introduced into *S. pyogenes* and the lox71-*erm*-lox66 cassette was removed. For the complementation of the Δ*yebC* mutant, the *yebC* gene with native promoter and terminator was cloned into the modified p7INT integrative vector^68^. Subsequently, the C-terminal 3x FLAG tag and mutations were introduced into the *yebC* sequence and the p7INT derivatives were introduced into the *S. pyogenes* genome by electroporation and antibiotic selection.

The reporter proteins for ribosome stalling were also introduced using the p7INT vector. The fusion protein 3x FLAG-sfGFP-mKate was cloned into p7INT and the mutations were introduced into sfGFP-mKate linker. The constructs were then introduced to *S. pyogenes* genome. The detailed mutagenesis protocols are provided in the Supplementary information.

### Orthogonal Organic Phase Purification (OOPS)

#### Cell culture and UV irradiation

*S. pyogenes* cells were grown in 30 mL of CDM until the mid-logarithmic growth phase. Each culture was grown in two batches: in one batch, CDM was supplemented with ^13^C^15^N-labelled L-lysine and L-arginine, in the other – with the non-labelled amino acids. The cells were rapidly cooled in an ice-water bath and collected by centrifugation for 5 min at 4500 g +4°C. The cells were re-suspended in 10 mL of ice-cold PBS and transferred to the cooled Petri dishes. One batch was UV irradiated with 600 mJ cm^-2^ UV 254 nm on the Bio-Link BLX cross-linker (Vilber), and the other batch was kept on ice. The UV+ and UV-batches were merged and the cells were collected by centrifugation for 5 min at 4500 g +4°C. The experiment was performed in four replicates: in two of the replicates, the UV+ culture was SILAC labelled, and in the other two replicates, the UV-culture was labelled.

#### Isolation of RBPs

The cell pellets were re-suspended in 400 µL of Disruption solution (50 mM Tris-HCl pH 7.6, 10% sucrose, 5 mM EDTA). The bacteria were transferred to screw-cap 2 mL tubes with 0.1 mm glass beads (Roth) and disrupted on Fastprep-24 5G (MP Biomedicals) using settings for *S. pyogenes* (speed 6.0, time 20 s). 20 µL of the lysates were collected and analysed by MS (Input samples). The rest of the lysates were subjected to phenol-chloroform extraction with 1 mL of Trizol and 0.2 mL of chloroform and the interfaces between the organic and the aqueous phases were collected as described in^4^. The interfaces were subjected to two more rounds of phenol-chloroform extraction. The third interfaces were precipitated by mixing with 900 µL of methanol and centrifugation for 10 min 14000 g at room temperature. The pellets were washed with 500 µL of methanol, dried for 10 min at the bench, dissolved by pipetting and vortexing in 100 µL of RBP buffer (100 mM TEAB, 1 mM MgCl_2_, 1% SDS) and stored at -20°C. On the next day, the samples were incubated at 95°C for 20 min and cooled to room temperature. 2 µL of RNase A/T1 mix (Thermo Scientific) was added and the samples were incubated for 2 h at 37°C. Additional 2 µL of RNase A/T1 mix was added and the samples were incubated overnight at 37°C. The samples were subjected to phenol-chloroform extraction with 0.5 mL of Trizol and 0.1 mL of chloroform. The organic phases were collected, mixed with 9 vol. of methanol and incubated at -20°C for 1 h. After centrifugation for 30 min at 20000 g +4°C, the supernatant was discarded and the precipitated proteins were gently washed with 500 µL of methanol and dried on the bench for 10 min. The proteins were solubilized in 40 µL of 1x Laemmli sample buffer without dye (60 mM This-HCl pH 6.8, 2% SDS, 10% glycerol, 12.5 mM EDTA, 150 mM β-mercaptoethanol) by incubation at 95°C for 10 min.

#### Mass spectrometry and data analysis

The samples were analysed on an Orbitrap Fusion Lumos (Thermo Scientific) that was coupled to a 3000 RSLCnano UPLC (Thermo Scientific). Detailed parameters for the mass spectrometry can be found in the Supplementary information. Raw files were processed with Proteome Discoverer 2.4 (Thermo Scientific) using SEQUEST HT for peptide identification and *S. pyogenes* UniProt FASTA protein database. MS1-based peptide ion quantification for SILAC was enabled. Peptide-spectrum-matches (PSMs) were filtered to a 1% FDR level using Percolator employing a target/decoy approach. The protein FDR was set to 1%. Further data processing was carried out in R and Perseus (v. 1.6.2.3). All contaminant proteins were filtered out. SILAC ratios were log2 transformed. In order to correct for mixing errors, the median SILAC ratio of the Input samples was determined and subtracted from the ratio of the corresponding OOPS samples. Proteins with a SILAC ratio count < 2 were removed. Next, quantile normalization was performed separately for each experiment (R package preprocessCore) and the resulting OOPS enrichment values were used for further analysis. Statistical significance of OOPS enrichment was assessed using the R package Limma^69^. Resulting p-values were corrected for multiple testing employing the approach by Benjamini-Hochberg^70^. Proteins with an adjusted *p*-value < 0.05 were considered to have statistically significant OOPS enrichment.

### RBS-ID

#### Cell culture and UV irradiation

*S. pyogenes* cells were grown in 500 mL of CDM with non-labelled L-Lysine and L-arginine until the mid-logarithmic growth phase. The cells were rapidly cooled on ice-water bath and collected by centrifugation for 5 min at 4500 g +4°C. The cells were re-suspended in 180 mL of ice-cold PBS, transferred to a cooled tray and UV irradiated with 600 mJ cm^-2^ UV-C on Bio-Link BLX cross-linker (Vilber). The cells were collected by centrifugation for 5 min at 4500 g +4°C.

#### Sample preparation

The cells were disrupted and three rounds of phenol-chloroform extraction were performed according to the OOPS protocol. After methanol precipitation, the third interface was dissolved in 1 mL of Reduction buffer (100 mM TEAB; 20 mM DTT) and incubated at room temperature for 1 h. Alkylation was performed with 40 mM 2-chloroacetamide for 2 h at 37°C. CaCl_2_ was supplemented to 1 mM concentration and protein fragmentation was performed with 0.5 µg of Trypsin and 0.5 µg of LysC overnight at 37°C. After adjusting the sample volume to 1.2 mL, it was mixed with 6 mL of buffer RLT supplemented with 40 mM DTT and 5.4 mL of 100% ethanol. The RNA-peptide conjugates were purified with RNeasy Midi kit (Qiagen) according to the manufacturer’s instructions, except that the centrifugation speed at binding was 500 g and RW1 buffer was omitted. 240 µL of the sample was mixed with 13 µL of 1 M Tris-HCl pH 7.6 and 1 mL of 48% hydrofluoric acid and incubated at 4°C overnight. The sample was dried with SpeedVac vacuum concentrator (Thermo Scientific) supplemented with CaCO_3_ trap and located under the fume hood.

#### Mass spectrometry and data analysis

The MS parameters can be found in the Supplementary information. In order to discover unknown modifications generated from UV crosslinking, an open search was performed using MSFragger (version 3.0) within FragPipe (version 13.0)^71^, using default settings (UniProt proteome database: *S. pyogenes*). Peptide matches were filtered to 1% FDR using PeptideProphet and Philosopher (version 3.2.9)^72^. The open search resulted in identification of three major XL modifications: uridine (+244.069536 Da), uridine –H_2_O (+226.05897 Da) and uridine –NH_3_ (+227.042987 Da). These modifications were selected as variable modifications (considering all amino acids) in a closed search using MSFragger, as described.

### Immunoprecipitation and PNK labelling of *S. pyogenes* RBPs

#### Cell culture and UV irradiation

*S. pyogenes* cells with the 3x FLAG tagged genes (Supplementary table 3) were grown in two batches of 25 mL of CDM with non-labelled L-Lysine and L-arginine until the mid-logarithmic growth phase. The cells were rapidly cooled on ice-water bath and collected by centrifugation for 5 min at 4500 g +4°C. The cells were re-suspended in 10 mL of ice-cold PBS and transferred to the cooled Petri dishes. The UV-sample was kept on ice and UV+ sample was irradiated with 600 mJ cm^-2^ UV-C on Bio-Link BLX cross-linker (Vilber). The cells were collected by centrifugation for 5 min at 4500 g +4°C.

#### Immunoprecipitation of the cross-linked protein-RNA complexes

The PNK assay was performed according to Holmqvist et al^26^. The cell pellets of UV- and UV+ samples were re-suspended in 800 µL of NP-T buffer (50 mM Na_2_HPO_4_ pH 8.0, 300 mM NaCl, 0.05% Tween 20) and bacteria were disrupted with 0.1 mm glass beads (Roth) on Fastprep-24 5G (MP Biomedicals) using the settings for *S. pyogenes* (speed 6.0, time 20 s). The lysates were cleared by centrifugation for 10 min at +4°C 10000 g and the supernatants were collected. For each sample, 15 µL of Anti-FLAG M2 Magnetic Beads (Sigma) were washed with NP-T using the DynaMag magnetic stand (Invitrogen) and re-dissolved in 100 µL of NP-T buffer. In a 2 mL tube, 500 µL of the cleared cell lysates, 1 mL of ice-cold NP-T buffer and 100 µL of the beads were mixed and rotated for 2 h at 4°C. The beads were collected by centrifugation for 1 min at 800 g +4°C, re-suspended in 200 µL of NP-T buffer and transferred to clean 1.5 mL tubes. The beads were washed once with 500 μL of High-salt buffer (50 mM Na_2_HPO_4_ pH 8.0, 1 M NaCl, 0.05% Tween 20) and twice with 500 μL of NP-T buffer. The beads were re-suspended in 100 µL of 1:1000 RNase I (10 U/μL; Thermo Scientific) dilution in NP-T, incubated for 3 min at 37°C with shaking at 1100 rpm and cooled for 2 min on ice. The beads were washed once with 500 μL of High-salt buffer and twice with 500 μL of CIAP/PNK wash buffer (50 mM Tris-HCl pH 7.4, 100 mM NaCl, 10 mM MgCl_2_). The RNA fragments were dephosphorylated with 10 U of CIAP (Thermo Scientific) at 37°C for 30 min with shaking at 1100 rpm. The beads were once again washed with High-salt buffer and twice with CIAP/PNK wash buffer. The magnetic beads of both UV- and UV+ samples were separated into two batches: PNK- and PNK+. The magnetic beads were mixed with 16 µL of mQ, 2 µL of 10x PNK buffer A, 1 µL of γ-32P ATP (10 µCi/µL; 3.3 µM; Hartmann Analytic) and incubated at 37°C for 30 min with 1100 rpm shaking without or with 1 µL of T4 PNK (10 U/µL; Thermo Scientific). The beads were washed with High-salt buffer and twice with CIAP/PNK wash buffer. Afterwards the beads were re-dissolved in 12 µL of 0.2 mg/mL 3x FLAG peptide (Sigma-Aldrich) in TBS and the cross-linked protein-RNA complexes were eluted by incubation at 37°C for 30 min 1100 rpm. The beads were magnetized, and the supernatants were collected.

#### Electrophoresis, phosphor imaging and Western blot

9 µL of the eluted protein-RNA complexes were resolved on NuPAGE 4-12% Bis-Tris Protein Gel with MOPS buffer (Thermo Scientific) and transferred to the 0.45 µm nitrocellulose membrane (Cytiva) using the semi-wet transfer apparatus (Thermo Scientific). The membrane was washed with PBS and incubated with the Phosphor screen for 24 h. The radioactive signal was detected using the Typhoon FLA 9500 imager (Cytiva). Western blotting was then performed using 1:5000 Monoclonal ANTI-FLAG M2 antibody produced in mouse (Sigma).

### Bioinformatic analysis of YebC proteins

The structure of *S. pyogenes* YebC was modelled on the basis of CBU_1566 from *Coxiella burnetii* (PDB 4F3Q) with SWISS-MODEL webserver^73^. The structure was visualized with PyMOL Molecular Graphics System, Version 3.0 Schrödinger, LLC. The electrostatics calculations were performed using the APBS software^74^. The amino acid sequences of *S. pyogenes* YebC (Spy_0316) and its homologs were downloaded from the KEGG database^20^. The sequences were aligned with Clustal Omega^75^. The alignment and secondary structures of Spy_0316 and *Mus musculus* TACOI (PDB 5EKZ) were visualized using ESPript^76^. The distribution of YebC_I and YebC_II clusters was extracted from the PANTHER database^77^ and visualized using the iTOL browser^78^.

### RNA sequencing and MS proteomics for *S. pyogenes* Δ*yebC* strain

#### Growth and collection of cells

The *WT*, Δ*yebC* and Δ*yebC / yebC+* strains (Supplementary table 3) were grown in THY until the mid-logarithmic (Log) and early stationary (Stat) growth phases. For RNA-seq 10 mL of cell cultures were mixed with 5 mL of Stop solution (5% phenol in absolute ethanol) and cooled in an ice-water bath. For MS proteomics 10 mL of cell cultures were cooled in an ice-water bath. The cells were precipitated by centrifugation for 5 min at 4500 g +4°C. The pellets were frozen and stored at -80°C. The experiment was performed in three independent replicates.

#### RNA sequencing and data analysis

The cell pellets were re-suspended in 500 µL of Disruption solution (50 mM Tris-HCl pH 7.6, 10% sucrose, 5 mM EDTA). The bacteria were transferred to the screw-cap 2 mL tubes with 0.1 mm glass beads (Roth) and disrupted on Fastprep-24 5G (MP Biomedicals) using the settings for *S. pyogenes* (speed 6.0, time 20 s). Phenol-chloroform extraction was performed with 1 mL of Trizol and 0.2 mL of chloroform and the aqueous phase was collected. One more round of extraction was performed with the equal volume of 1:1 acidic phenol-chloroform mixture and the residual amounts of phenol were removed by chloroform extraction. The RNA was precipitated by mixing the aqueous phases with an equal volume of 2-propanol, incubating for 1 h at -20°C and centrifuging for 30 min at 20000 g +4°C. The pellets were washed with 500 µL of 80% ethanol, dried on the bench and dissolved in 100 µL of mQ water. The cDNA libraries were prepared according to the protocol provided in the Supplementary information. Sequencing of the libraries was performed on NovaSeq 6000 (Illumina) in a 2×100 bp paired-end mode.

For the analysis of RNA-seq data, reads were filtered with a minimum quality score of 10 and a length of at least 18 nt, cleaned from adapter sequences with the parameters: ‘-a AGATCGGAAGAGCACACGTCTGAACTCCAGTCA-AAGATCGGAAGAGCGTCGTGTAGGGAAAGAGTGT’ using Cutadapt (v4.4)^79^ and mapped against the reference genome NC_002737.2 using STAR (v2.7.3a)^80^ in ‘random best’ and ‘end-to-end’ modes. BAM files were sorted and indexed using Samtools (v1.19.2)^81^, and PCR duplication artefacts were removed using UMI Tools (v1.1.5)^82^. Gene counts of annotated RefSeq genes were determined with featureCounts (v2.0.1)^83^ and differentially expressed genes (DEG) were calculated with three replicates per condition and growth phase using DESeq2 (v1.38.0)^84^, and a threshold of adjusted *p*-value < 0.05 and absolute log2 fold change > 1 was used to call DEGs. We performed the aforementioned analysis steps using a customized pipeline implemented in Snakemake (v8.4.8), a workflow management system^85^.

#### MS proteomics and data analysis

The cell pellets were re-suspended in 800 µL of Lysis buffer (20 mM Tris-HCl pH 8.0, 100 mM NH_4_Cl, 10 mM MgCl_2_, 0.4% Triton X-100, 0.1% NP-40). The bacteria were transferred to 2 mL screw-cap tubes with 0.1 mm glass beads (Roth) and disrupted on Fastprep-24 5G (MP Biomedicals) using the settings for *S. pyogenes* (speed 6.0, time 20 s). The lysates were cleared by centrifugation and analysed by MS. The MS parameters can be found in the Supplementary information. Raw data analysis was performed using Spectronaut (Biognosys AG, Zurich, Switzerland) version 17 in directDIA+ deep mode with reviewed UniProt databases (Streptococcus pyogenes serotype M1 – version 2024_02_05, 1,690 canonical entries). Methionine oxidation and Acetyl (Protein N-term) were set as a variable, and carbamidomethylation on cysteine residues was used as a static modification. The FDR for PSM-, peptide-, and protein-level was set to 0.01. All tolerances were set to dynamic for pulsar searches. All downstream data processing was carried out using R and R studio. Extracted features were exported from Spectronaut for statistical analysis with MSstats^86^ using default settings. Briefly, for each protein, features were log-transformed and fitted to a mixed-effect linear regression model for each sample. The model estimated fold change and statistical significance for all compared conditions. The Benjamini–Hochberg method accounted for multiple tests, and a *p*-value adjustment was performed on all proteins that met the fold-change cut-off. Significantly different proteins were determined using a threshold of fold-change greater than two and an adjusted *p*-value < 0.05.

### Infection of human macrophages with *S. pyogenes*

#### Cell culture

For the generation of primary human macrophages, PBMCs were purified from buffy coats derived from the German Red Cross Blood Transfusion Service. Briefly, buffy coats were diluted 1:1 in PBS and separated in Pancoll density gradients (1.077 g/ml, Pan Biotech; 800 g, 30 min, room temperature, without break). PBMCs were washed twice with PBS and plated at a density of 10x10^7^ cells in RPMI in 10 cm tissue culture dishes (Sarstedt) for 3h. Then the medium was replaced by RPMI supplemented with 10% human serum type AB and 100ng/mL GM-CSF (Miltenyi Biotec). The following day, cells were washed once with PBS and further incubated in RPMI supplemented with 10% human serum type AB and 100ng/mL GM-CSF for 5 days in a humidified atmosphere containing 5% CO_2_. Media was replaced for RPMI + 10% human serum, type AB without GM-CSF and macrophages were incubated for a further day before being plated in 24-well plates at a density of 2×10^5^ cells per well.

#### In vitro infection of primary macrophages

Infection assays were performed with *S. pyogenes* from mid-logarithmic phase cultures in THY (OD_600 nm_: ∼0.5). Bacteria were centrifuged (4700 g, 5 min), washed with PBS and passed ten times through a 27 Gauge syringe insert in a 1 ml syringe (Braun) to separate the cocci chains. Cells were infected for 1h with a Multiplicity of Infection (MOI) of 5 in RPMI + 1 % human serum including centrifugation for 5 min at 300 g for synchronization. For gentamicin protection assays, macrophages were subsequently incubated with RPMI + 10% human serum and 100 μg/mL Gentamicin for 1h to eliminate extracellular bacteria. Macrophages were further incubated with 10μg/mL Gentamicin until the end of the experiment. To determine intracellular bacterial loads, macrophages were lysed at the indicated time points using 0,1% Saponin in PBS, and serial dilutions of the lysates were plated on THY agar plates incubated for 24h at 37°C with 5% CO_2_. For LDH and ELISA experiments, following the 1h infection period, macrophages were incubated with RPMI + 10% human serum and 1% penicillin/streptomycin. Supernatants were harvested at 4 h and 24 h post-infection.

#### ELISA and LDH assays

Cytokines and LDH in macrophage supernatants were measured by R&D Systems DuoSet ELISA Development Systems (human IL-1β, DY201; human IL18, DY318; human IL-6, DY206; human TNF, DY210; human IL-8, DY208) or using the LDH Assay Kit (Abcam) according to the manufacturer’s instructions. The percentage of cytotoxicity was calculated as follows:

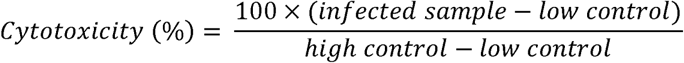

### iCLIP for YebC in *S. pyogenes*

#### Cell culture and UV irradiation

*S. pyogenes WT* and *yebC:3xFLAG* strains were grown in 400 mL of CDM with non-labelled L-Lysine and L-arginine until the mid-logarithmic growth phase. The cells were rapidly cooled in an ice-water bath and collected by centrifugation for 5 min at 4500 g +4°C. The cells were re-suspended in 100 mL of ice-cold PBS, transferred to cooled trays and UV irradiated with 600 mJ cm^-2^ UV-C on Bio-Link BLX cross-linker (Vilber). The cells were collected by centrifugation for 5 min at 4500 g +4°C. The UV-irradiated *WT* cells represent the “WT_UV+” control. Growth and UV irradiation of *yebC:3xFLAG* cultures were performed in two biological replicates. An additional *yebC:3xFLAG* culture was not treated with UV-C and represents the “YebC_UV-“ control.

#### Immunoprecipitation of the cross-linked YebC-RNA complex

The iCLIP protocol is based on^33^ with bacteria-specific modifications adapted from^87^. For each sample the cell pellets were re-suspended in 4×800 µL of NP-T buffer (50 mM Na_2_HPO_4_ pH 8.0, 300 mM NaCl, 0.05% Tween 20) and bacteria were disrupted with 0.1 mm glass beads (Roth) on Fastprep-24 5G (MP Biomedicals) using the settings for *S. pyogenes* (speed 6.0, time 20 s). The lysates were cleared by centrifugation for 10 min at +4°C 10000 g and the supernatants were collected. For each sample 4×550 µL of cell lysates were collected and combined. The protein concentrations were measured and adjusted to 1 mg/mL with NP-T buffer. 1:25000 dilution of RNase I (10 U/μL; Thermo Scientific) was added and the lysates were incubated at 37°C with 1100 rpm shaking for 3 min and on ice for 2 min. At this step 15 µL of UV-irradiated lysates of *yebC:3xFLAG* strain were collected and served as “size-matched input” (SMI) controls.

For each sample, 15 µL of Anti-FLAG M2 Magnetic Beads (Sigma) were washed with NP-T using the DynaMag magnetic stand (Invitrogen) and re-dissolved in 200 µL of NP-T buffer. 2.2 mL of lysates were mixed with 20 mL of NP-T buffer and the washed magnetic beads, and incubated with rotation at 4°C for 2 h. The beads were collected by centrifugation for 1 min at 800 g +4°C, re-suspended in 200 µL of NP-T buffer and transferred to clean 1.5 mL tubes. The beads were washed once with 500 μL of High-salt buffer (50 mM Na_2_HPO_4_ pH 8.0, 1 M NaCl, 0.05% Tween 20) and twice with 500 μL of MES-PNK buffer (25 mM MES pH 6.0, 50 mM NaCl, 10 mM MgCl_2_, 0.1% Tween 20). The beads were then re-dissolved in 20 µL of MES-PNK buffer with 10 mM DTT and 25U of T4 PNK (Thermo Scientific) and the dephosphorylation reaction was performed for 30 min at 37°C with shaking at 1100 rpm. The beads were once again washed with High-salt buffer and twice with CIAP/PNK wash buffer (50 mM Tris-HCl pH 7.4, 100 mM NaCl, 10 mM MgCl_2_). The magnetic beads were mixed with 16 µL of mQ, 2 µL of 10x PNK buffer A, 1 µL of γ-32P ATP (10 µCi/µL; 3.3 µM; Hartmann Analytic) and 1 µL of T4 PNK and incubated at 37°C for 30 min with shaking at 1100 rpm. The beads were washed with High-salt buffer and twice with CIAP/PNK wash buffer. The beads were then re-dissolved in 12 µL of 0.2 mg/mL 3x FLAG peptide (Sigma-Aldrich) in TBS and the cross-linked YebC-RNA complexes were eluted by incubation at 37° for 30 min with shaking at 1100 rpm. The beads were magnetized and the supernatants were collected.

#### Sequencing of RNA molecules cross-linked to YebC

The YebC_UV+ samples and controls (WT_UV+, YebC_UV- and SMI) were resolved on NuPAGE 4-12% Bis-Tris Protein Gel with MOPS buffer (Thermo Scientific). The cross-linked YebC-RNA complexes were transferred to the 0.45 µm NC membrane using the semi-wet transfer apparatus (Thermo Scientific). The membrane was washed with PBS and incubated with the Phosphor screen for 2 h. The radioactive signal was detected using the Typhoon FLA 9500 imager (Cytiva). The zones corresponding to the labeled RNA fragments in YebC_UV+ samples (∼30-65 kDa) were excised from the membrane using the phosphor imaging print-out as a mask. The pieces of nitrocellulose membrane were submerged in 150 µL of PK buffer (100 mM Tris-HCl pH 7.4, 50 mM NaCl, 10 mM EDTA) and treated with 10 µL of 20 mg/mL proteinase K, RNA grade (Thermo Scientific) at 37°C shaking at 1100 rpm for 20 min. 150 µl of PK buffer with 7 M urea was added and the samples were incubated for a further 20 min at 37 °C and 1100 rpm. The solutions were collected and the RNA was extracted with equal volumes of 1:1 acid phenol-chloroform. The aqueous layers were collected and residual phenol was removed by chloroform extraction. The RNA was ethanol precipitated with GlycoBlue Coprecipitant (Thermo Scientific) and re-dissolved in 11 µL of mQ. The cDNA libraries were prepared according to the protocol provided in the Supplementary information. Sequencing of the libraries was performed on the NovaSeq 6000 (Illumina) in a 2×100 bp paired-end mode.

#### Analysis of iCLIP data

iCLIP reads were pre-processed as described in Bush et al.^35^. In brief, reads with a high quality (minimum quality score of 20) in the barcode region were filtered using FASTX-Toolkit (v0.0.13) [http://hannonlab.cshl.edu/fastx_toolkit/] and seqtk subseq (v1.3) [https://github.com/lh3/seqtk], demultiplexed and trimmed using Flexbar (v3.5.0)^88^. Reads with a length of at least 18 nt were then mapped against the reference genome NC_002737.2 using STAR (v2.7.10a_alpha_220314) in ‘random best’ and ‘Extend5pOfRead1’ modes. The BAM files were sorted and indexed using Samtools (v1.9), and PCR duplication artefacts were removed using UMI Tools (v1.1.0).

For each sample transcriptional coverage was calculated with custom R scripts using the GenomicRanges package^89^. Nucleotides located directly upstream of the mapped reads were considered cross-linking sites and for each genomic position the number of the cross-linking sites was calculated. The coverage and the number of the cross-linking sites at each genomic position were divided by the total number of mapped reads and multiplied by 1×10^6^ to obtain the normalized transcripts per million (TPM) values. The normalized coverage and the density of cross-linking sites of six *S. pyogenes* rRNA operons were collapsed to the first operon (SPY_RS00070, SPY_RS00080 and SPY_RS00085). Using a sliding window approach, the peaks of the cross-linking sites were mapped: these are 30 nt long regions where the sum of TPMs for the cross-linking sites is greater than three and more than five times greater than that of the 30 nt long up- and downstream regions.

### Association of YebC with *S. pyogenes* ribosomes

#### Growth, collection and disruption of cells

The *S. pyogenes* strain *yebC:3xFLAG* was grown in 150 mL of THY until the mid-logarithmic (Log) and stationary (Stat) growth phases. The cells were collected by rapid filtration using a vacuum funnel (Merck) with 0.22 µm filter, immediately frozen in liquid nitrogen and stored at -80°C. The 10 mL stainless steel mill jars (Retsch) were cooled in liquid nitrogen. The frozen cell pellets were transferred to the cooled mill jars along with 2.5 mL of Lysis buffer (20 mM Tris-HCl pH 8.0, 100 mM NH_4_Cl, 10 mM MgCl_2_, 0.4% Triton X-100, 0.1% NP-40, 1 mM chloramphenicol, 100 U/mL DNase I). The cells were disrupted on MM400 swing mill (Retsch) with five rounds of grinding for 1 min at 30 Hz. Between the grinding rounds the mill jars were cooled in liquid nitrogen. The cell powders were collected and melted on ice. The lysates were centrifuged at 20000 g +4°C for 20 min and the supernatants were collected.

#### Sucrose density gradient centrifugation and collection of fractions

The 10% and 50% sucrose buffers (20 mM Tris-HCl pH 8.0, 100 mM NH_4_Cl, 10 mM MgCl_2_, 0.5 mM DTT, sucrose) were prepared and cooled on ice. The 10-50% sucrose gradients (vol. = 12 mL) were formed in SW40 ultracentrifuge tubes (Beckman Coulter) using Gradient Station (Biocomp). 0.5 mL of cell lysates were applied on top of the gradients and the tubes were centrifuged in SW 40 Ti Swinging-Bucket Rotor (Beckman Coulter) at 35000 rpm for 2 h at 4 °C. The gradients were fractionated with Piston Gradient Fractionator (Biocomp) equipped with FC 203B Fraction Collector (Gilson). 15 × 0.8 mL fractions were collected. During the fractionation, OD_260 nm_ measurements were performed using the Triax Flow Cell (Biocomp).

#### Analysis of fractions

500 µL of fractions 1 to 12 were mixed with 125 µL of trichloroacetic acid, incubated on ice for 1 h and centrifuged for 5 min at 14000 g +4°C. The pellets were washed twice with acetone and incubated at 65°C for 10 min. The pellets were dissolved in 20 µL of Laemmli loading buffer and incubated at 95°C for 10 min. The samples were resolved on two Laemmli SDS-PAGE gels. One of the gels was stained with Coomassie. Proteins from the other gel were semi-dry transferred to a 0.45 µm NC membrane and western blotting was performed using 1:5000 dilution of Monoclonal ANTI-FLAG M2 antibody produced in mouse (Sigma).

### Ribosome profiling for *S. pyogenes yebC* mutants

#### Cell culture and isolation of monosomes

The ribosome profiling protocol is based on Galmozzi et al^90^ and was adapted to *S. pyogenes*. The *WT*, Δ*yebC*, Δ*yebC / yebC+* and Δ*yebC / yebC_M2+* strains were grown in 200 mL of THY until the mid-logarithmic growth phase. The experiment was performed in three biological replicates. The cells were collected and disrupted in the same way as for the study of the association of YebC with ribosomes. The lysates were centrifuged at 20000 g +4°C for 20 min and the supernatants were collected. The lysates were supplemented with 5 mM CaCl_2_ and treated with 0.1 U/OD_620 nm_ of Nuclease S7 (Merck) at 25°C with shaking at 400 rpm for 1 h. The reactions were stopped by adding EGTA to a concentration of 6 mM. 400 µL of the lysates were resolved on 10-50% sucrose density gradients in the same way as for the study of the association of YebC with ribosomes and the monosome fractions were collected.

#### Isolation and sequencing of the ribosome protected mRNA fragments

700 µL of the monosome fractions were mixed with 40 µL of 20% SDS and 750 µL of pre-warmed acid phenol-chloroform mix and incubated at 65 °C for 5 min with shaking at 1400 rpm. The samples were cooled on ice for 5 min and centrifuged at 20000 g +25°C for 2 min. The aqueous phases were collected and extractions were repeated with acid phenol-chloroform and chloroform. The RNA was then ethanol precipitated with GlycoBlue Coprecipitant (Thermo Scientific) and re-dissolved in 20 µL of mQ. For each sample 15 µg of RNA was resolved on 15% Urea TBE PAGE along with 15 and 45 nt control RNAs. Using the control RNA as guides the 15-45 nt long fragments were excised and isolated from the gel. The RNA was precipitated with 2-propanol and GlycoBlue Coprecipitant and re-dissolved in 15.5 µL of mQ. The cDNA libraries were prepared according to the protocol provided in the Supplementary information. The libraries were sequenced on NovaSeq 6000 (Illumina) in 100 bp single-end mode.

#### Analysis of ribosome profiling data

To obtain ribosome profiling footprints, reads were filtered for a minimum quality score of 10 and a length between 22 to 52 nt, cleaned from adapter sequence (3’end adapter: ‘ATCGTAGATCGGAAGAGCACACGTCTGAA’) using Cutadapt (v4.4) and mapped against the reference genome NC_002737.2 using STAR (v2.7.3a) in ‘random best’ and ‘end-to-end’ modes. BAM files were sorted and indexed using Samtools (v1.6). Random unique molecular identifiers that were introduced at the 5′ end (2 nt) and 3′ end (5 nt) of the footprint were detected and PCR de-duplication was performed using UMI Tools (v1.1.4). Reads mapping to rRNA or tRNA features as defined in the NCBI RefSeq annotation of NC_002737.2 were removed using Bedtools (v2.31.0)^91^. To obtain these pre-processed ribosome reads, we ran our customized ribo-seq snakemake pipeline up to the ‘filter_bam’ module. The source code for the ribo-seq pipeline is available at https://github.com/MPUSP/snakemake-bacterial-riboseq.

For all subsequent analyses, we selected reads having a length between 27 and 40 nt. The position of the ribosomal P site was mapped 15 nt upstream of the 3′ ends of the reads^38, 39^ and for each codon of *S. pyogenes* proteins, the number of reads representing the ribosomal A site was calculated using custom R scripts. To account for the difference in mRNA abundance, this number was divided by the sum of the numbers for the coding sequence. The resulting values were termed “pause scores” and reflect the occupancy of each codon by the translating ribosome. To identify codons for which pausing depends on the presence of YebC, six replicates of YebC+ samples (*WT* and Δ*yebC / yebC+* strains) were compared with six replicates of YebC-samples (Δ*yebC* and Δ*yebC / yebC_M2+*). To estimate the statistical significance of the difference, we used the edgeR software^92^, treating each codon as a gene and each coding sequence as a library. Pausing at the codons having an FDR adjusted *p*-value less than 0.001, a difference of more than 4-fold and an average pausing score in the up-regulated sample greater than 2 was considered to be affected by the mutation of YebC protein.

### Ribosome stalling reporter in *S. pyogenes*

The reporter strains (Supplementary table 3) were grown in 15 mL of THY until the mid-logarithmic growth phase. Expression of the reporter protein was induced by addition of 10 ng/mL anhydrotetracycline and the cells were grown for 30 min. The cell cultures were cooled in an ice-water bath and the cells were collected by centrifugation for 5 min at 4500 g +4°C. The cell pellets were re-suspended in 800 µL of NP-T buffer (50 mM Na_2_HPO_4_ pH 8.0, 300 mM NaCl, 0.05% Tween 20) and bacteria were disrupted with 0.1 mm glass beads (Roth) on Fastprep-24 5G (MP Biomedicals) using the settings for *S. pyogenes* (speed 6.0, time 20 s). The lysates were cleared by centrifugation for 10 min at +4°C 10000 g and the supernatants were collected. 5 µL of the lysates were resolved on a NuPAGE 4-12% Bis-Tris Protein Gel with MOPS buffer (Thermo Scientific) and transferred to a 0.45 µm nitrocellulose membrane (Cytiva) using the semi-wet transfer apparatus (Thermo Scientific). Western blotting was performed using 1:5000 dilution of Monoclonal ANTI-FLAG M2 antibody produced in mouse (Sigma). The expression of full-length fusion proteins was measured by densitometry of western blot photograph with Fiji^93^ and for each strain was normalized to the expression of P0 variant. The experiment was performed in three biological replicates and the statistical significance of the difference in protein expression relative to the P0 variant was estimated using the one-sample Student’s *t*-test.

### *In vitro* ribosome stalling reporter

The coding sequences of *E. coli yebC* paralogs *b1983* and *b1864* were amplified from genomic DNA of BW25113 strain and cloned into the pET-21a(+) plasmid. The proteins were expressed and purified according to the protocol provided in the Supplementary information. The templates for *in vitro* translation were designed on the basis of the DHFR control plasmid from the PUREexpress kit (New England Biolabs). The FLAG tag sequence and mutations were introduced to the *folA* N-terminus by two rounds of the whole-plasmid PCR with primers listed in Supplementary table 1, and circularization with KLD mix. The DHFR plasmids with the mutant versions of *folA* were linearized with the NotI FD restriction enzyme (Thermo Scientific) and used as templates for *in vitro* transcription using the MEGAscript T7 Transcription Kit (Thermo Scientific). The synthesized mRNAs were purified with the RNeasy Mini kit (Qiagen). *In vitro* translation was performed with the PURExpress in vitro protein synthesis kit (New England Biolabs). The 10 µL PURExpress reactions with or without 2 µM YebC variants were preincubated at 37°C for 5 min. mRNA templates were heated at 70°C for 1 min, cooled on ice and added to the reactions to a concentration of 1 µM. The reactions proceeded for 1 h at 37°C. Next, 0.2 mg/mL RNase A (Qiagen) was added to some reactions and the RNA was degraded for 10 min at 37°C. 2 µL of the reactions were resolved on a NuPAGE 4-12% Bis-Tris Protein Gel with MES buffer (Thermo Scientific) and transferred to a 0.2 µm nitrocellulose membrane (Cytiva) using the semi-wet transfer apparatus (Thermo Scientific). Western blotting was performed with 1:5000 of Monoclonal ANTI-FLAG M2 antibody produced in mouse (Sigma).

## Acknowledgements

We thank Thomas Wulff for providing sfGFP-mKate fusion, and Karin Hahnke, Florian Kondrot and Matthias Münzner for technical support. We are also grateful to Kürşad Turgay, Marc Erhardt and Rainer Nikolay for scientific discussions and critical reading of the manuscript. This work was funded by the Max Planck Society (for E.C.) and the German Research Foundation (DFG; Leibniz Prize for E.C.).

## Author contributions

D.I: Conceptualization, Methodology, Investigation, Formal Analysis, Software, Visualization, Writing – Original Draft. V.S.: Resources, Methodology, Investigation. R.A.: Data Curation, Formal Analysis, Software, Visualization, Writing – review & editing. K.A.: Methodology, Investigation, Formal Analysis. C.K.F: Methodology, Investigation, Formal Analysis. C.W.: Resources, Investigation. K.K.: Resources, Investigation, Formal Analysis. E.C.: Conceptualization, Funding Acquisition, Project Administration, Writing – Review & Editing.

## Data availability

All next generation sequencing data have been deposited in the European Nucleotide Archive (ENA) under the accession PRJEB78417. The proteomics mass spectrometry data have been deposited to the ProteomeXchange Consortium via the PRIDE partner repository with the dataset identifier PXD054642.

