## Supplementary information for "Novel RNA-binding protein YebC enhances translation of proline-rich amino acid stretches in bacteria"

### Supplementary figures

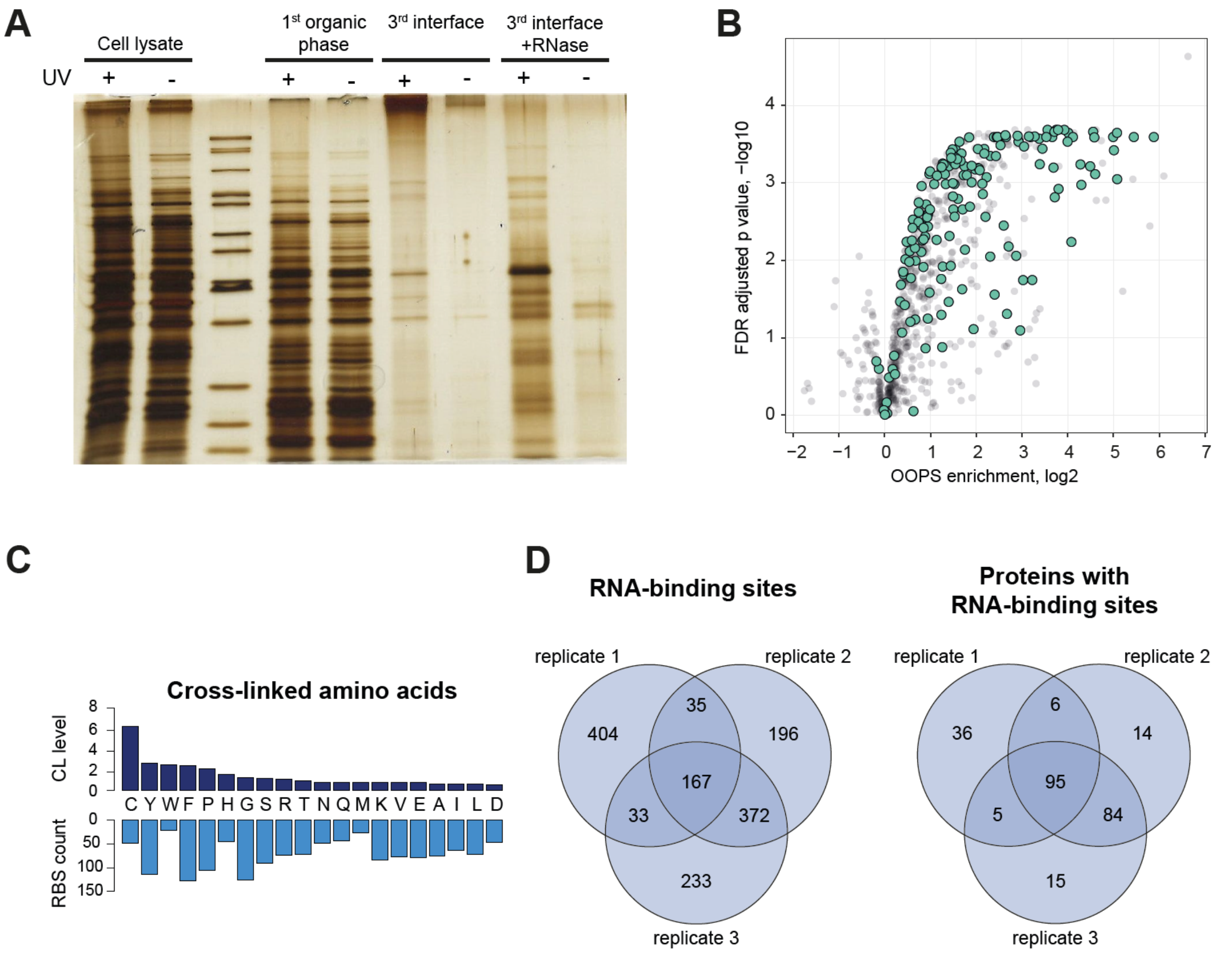


**Supplementary figure 1. Results of OOPS and RBS-ID for *S. pyogenes*.**

**(A)** UV-irradiated and control samples purified according to the OOPS protocol. *S. pyogenes* cell lysates were subjected to three rounds of acid guanidinium thiocyanate-phenol-chloroform phases partition, where the interfaces between the aqueous and organic phases were collected. The third interface was treated with RNase. The fractions representing the different steps of the protocol were resolved on SDS-PAGE and the gel was stained with silver.

**(B)** Volcano plot of OOPS enrichment values for *S. pyogenes* proteins. The annotated RBPs are depicted in green.

**(C)** Representation of amino acids at RNA-binding sites. The cross-linking levels were calculated by dividing the proportion of each amino acid in the identified RNA-binding sites by their proportion in the sequences of proteins where RNA-binding sites were identified. The corresponding counts of RNA-binding sites are shown below.

**(D)** Venn diagrams showing the numbers of RNA-binding sites and proteins with RNA-binding sites identified in three RBS-ID replicates and the overlap between them.

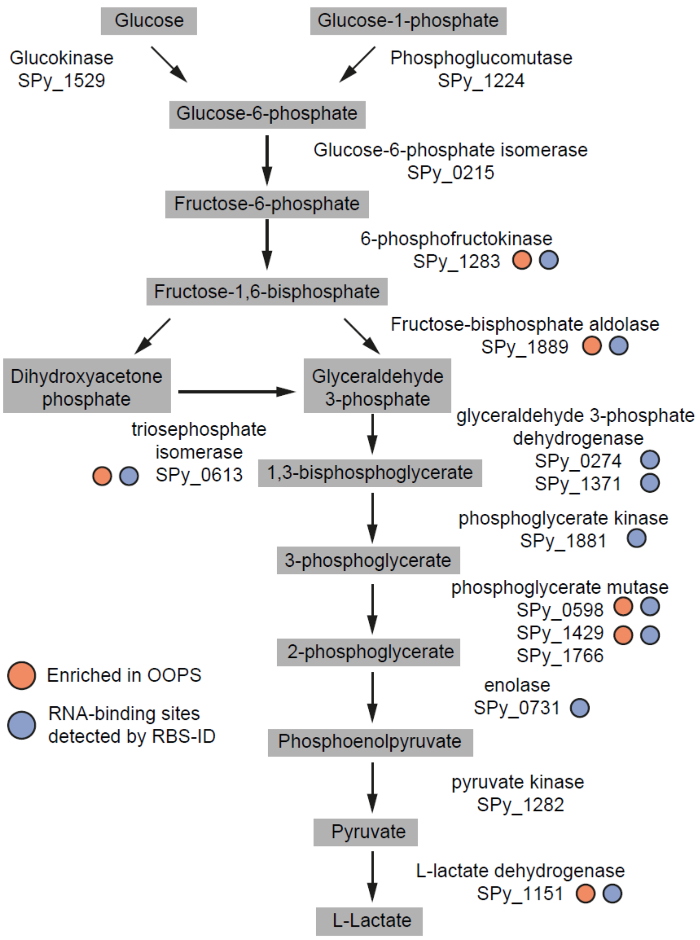


**Supplementary figure 2. OOPS enrichment and identification of RNA-binding sites for glycolysis enzymes.**

The genes encoding glycolysis enzymes were annotated according to KEGG database. The statistically significant OOPS enrichment and the detection of RNA-binding sites are indicated.

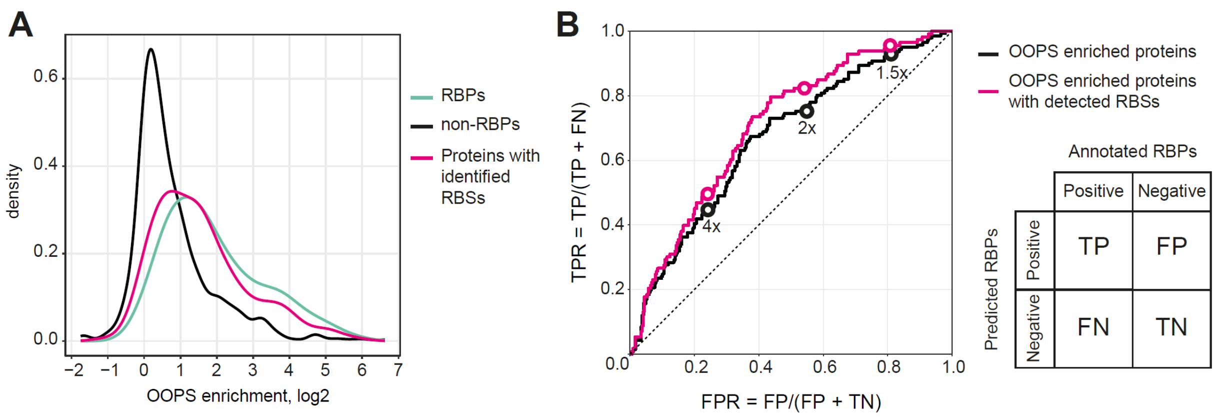


**Supplementary figure 3. Identification of candidates for novel RBPs.**

**(A)** Distribution of OOPS enrichment values for the annotated RBPs, non-RBPs and proteins with identified RNA-binding sites.

**(B)** ROC curve for the prediction of RBPs among the proteins with statistically significant OOPS enrichment. The annotated RBPs were considered true positives and all other proteins – true negatives. The OOPS enrichment values and detection of RNA-binding sites were used for classification. The OOPS enrichment cut-offs of 1.5, 2 and 4 are indicated on the plot.

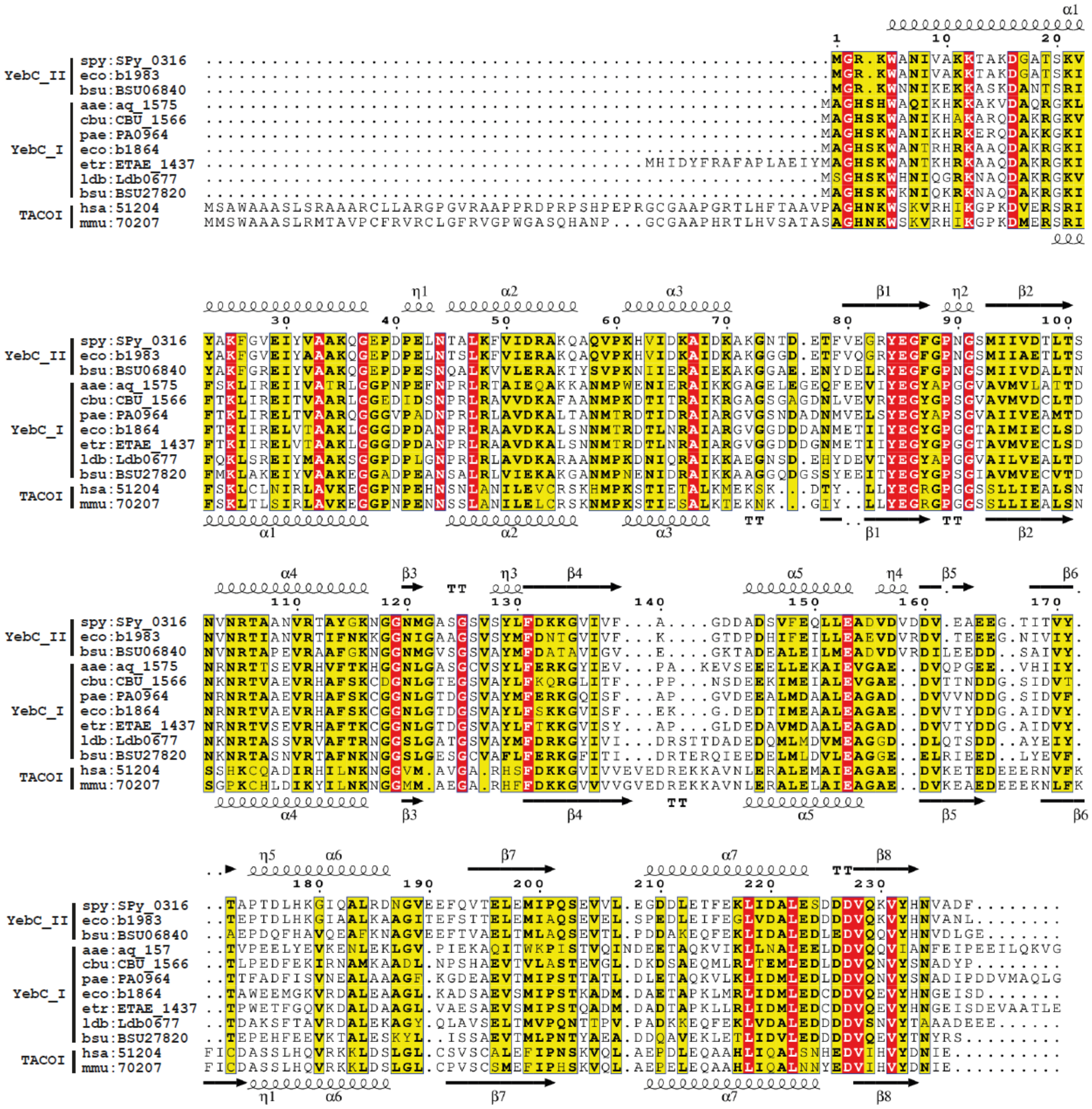


**Supplementary figure 4. Alignment of YebC homologs from different organisms.**
Amino acid sequences of YebC homologs from *S. pyogenes* (spy), *E. coli* (eco), *B. subtilis* (bsu), *Aquifex aeolicus* (aae), *Coxiella burnetii* (cbu), *Pseudomonas aeruginosa* (pae), *Edwardsiella piscicida* (etr), *Lactobacillus delbrueckii* (ldb), *Homo sapiens* (hsa) and *Mus musculus* (mmu) were aligned with Clustal Omega. Similar residues are written with black bold characters and boxed in yellow. Strictly conserved residues are boxed in red. Secondary structures of *S. pyogenes* YebC and *M. musculus* TACOI (PDB 5EKZ) are shown above and below the alignment respectively.

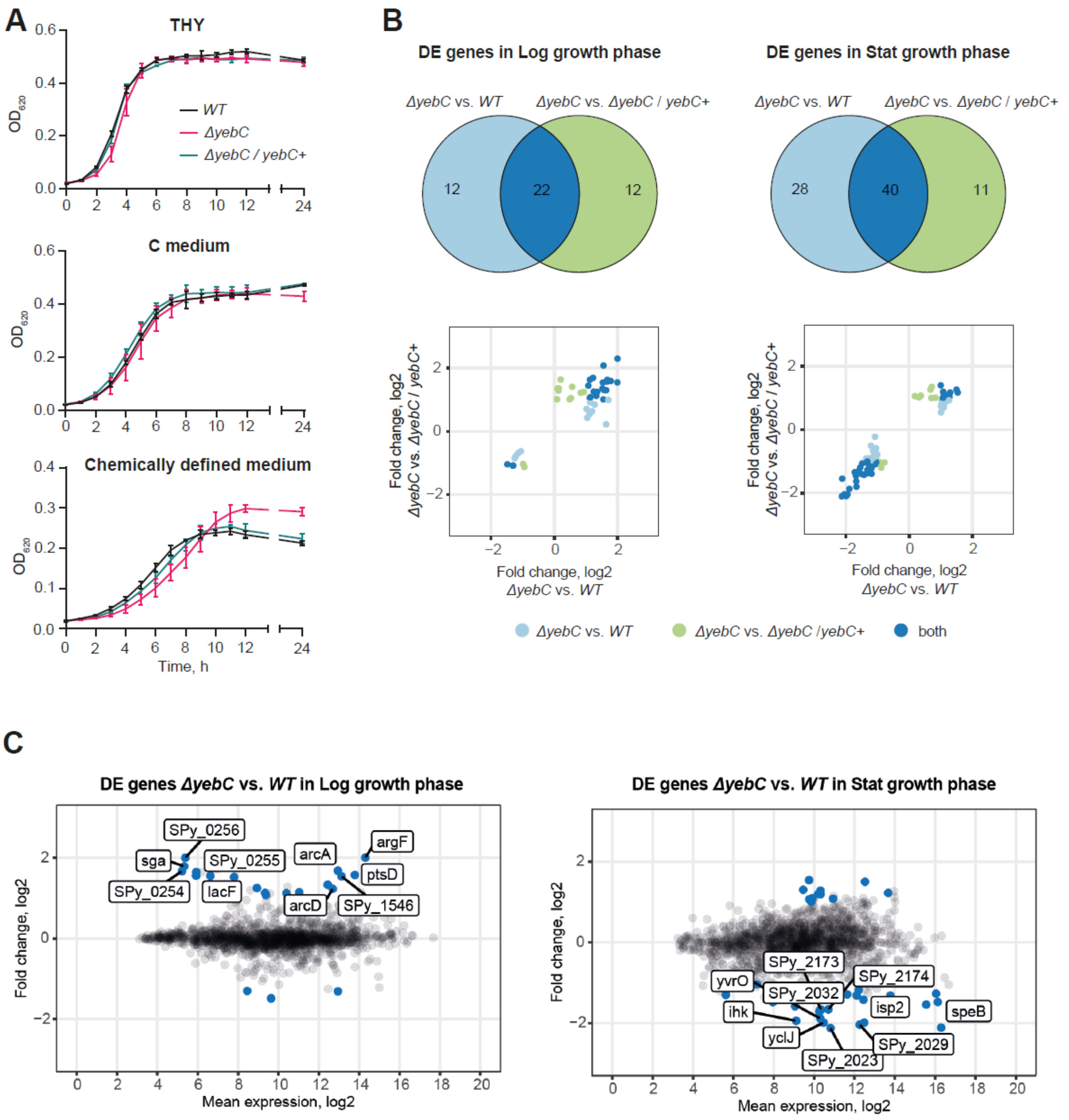


**Supplementary figure 5. Growth and transcriptome of *S. pyogenes ΔyebC* strain.**
**(A)** Growth curves of the *ΔyebC* and *ΔyebC / yebC+* strains in THY, C medium and CDM.
**(B)** Gene expression changes in the *ΔyebC* vs. *WT* and *ΔyebC* vs. *ΔyebC / yebC+* strains measured in mid-logarithmic (Log) and stationary (Stat) growth phases. The Venn diagrams show the numbers of differentially expressed (DE) genes identified uniquely identified in one or both of the comparisons described. The scatterplots below present the fold change of the DE genes in the comparisons. The colour indicates whether the DE genes were identified only in one or in both of the comparisons.
(C) MA plots indicating DE genes for the *ΔyebC* vs. *WT* comparisons in Log and Stat growth phases. Ten DE genes with the largest fold changes are indicated.

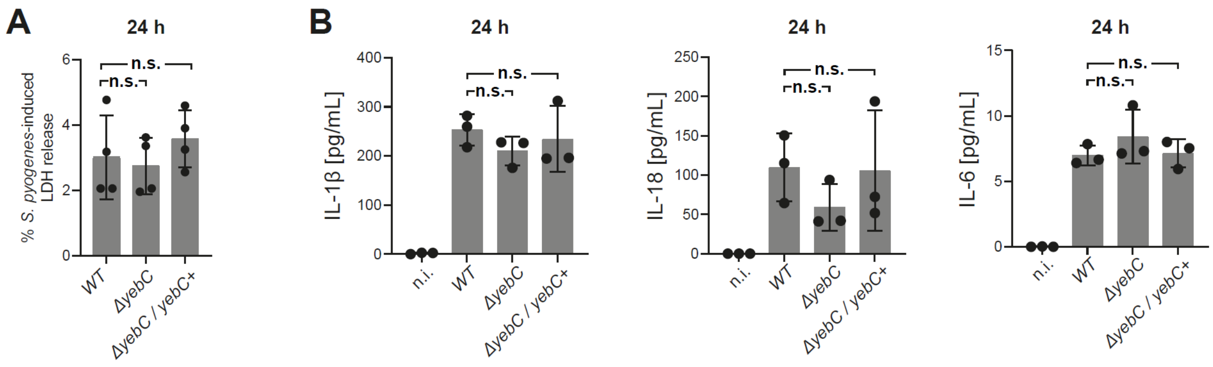


**Supplementary figure 6. Infection of macrophages.**

**(A)** LDH released by human macrophages infected with the WT, *ΔyebC*, and *ΔyebC / yebC*+ strains at 24 h (MOI 5:1). Bars represent the mean ± standard deviation (SD) of three biological replicates. One-way ANOVA with Holm-Šídák correction for multiple comparisons was applied for statistical analyses.

**(B)** IL-1β, IL-18, and IL-6 released by human macrophages infected with the *WT*, *ΔyebC*, and *ΔyebC / yebC*+ strains at 24 h (MOI 5:1). Bars represent the mean ± standard deviation (SD) of three biological replicates. One-way ANOVA with Holm-Šídák correction for multiple comparisons was applied for statistical analyses.

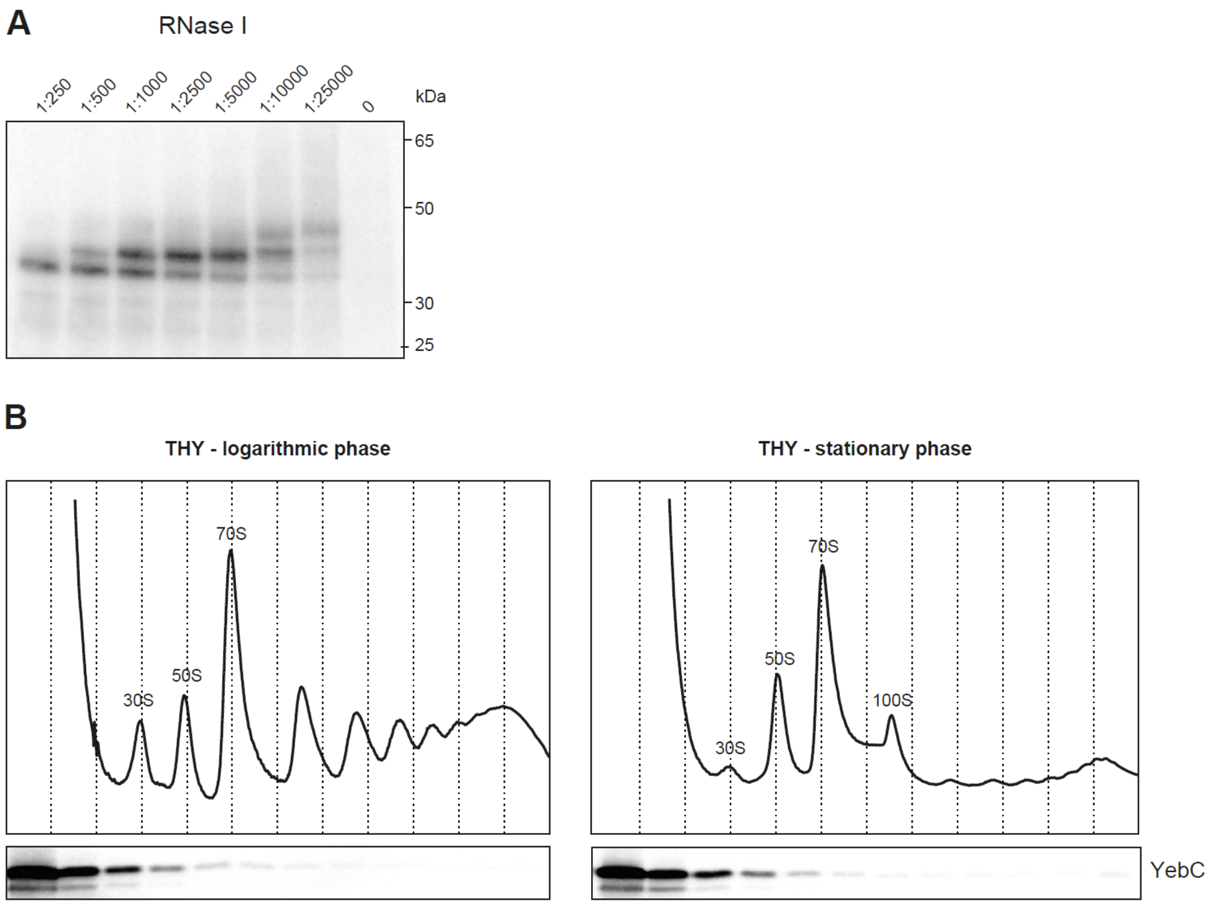


**Supplementary figure 7. Interaction of YebC with the ribosome.**
**(A)** Migration pattern of the cross-linked YebC-RNA complexes after treatment with RNase I. The complexes were immunoprecipitated using the anti-FLAG antibodies and treated with different dilutions of RNase I. After radioactive labelling of the RNA, the complexes were resolved on Bis-Tris gel and transferred to the nitrocellulose membrane. The radioactive signal is not visible in the absence of RNase I suggesting that the cross-linked RNA was too long to enter the gel. The mass of the complexes increases upon RNase I dilution and at dilutions greater than 1:10000 the complexes are represented by three distinct species.
**(B)** Association of YebC with the ribosome in the logarithmic and stationary growth phases. The *yebC:3xFLAG* strain was grown in THY medium until the logarithmic and stationary growth phases and rapidly collected. The lysates from each strain were resolved by sucrose density gradient ultracentrifugation. Sucrose density gradient fractions for each strain were probed with an anti-FLAG antibody. The results of the western blots are aligned with the respective $OD_{260\ nm}$ traces.

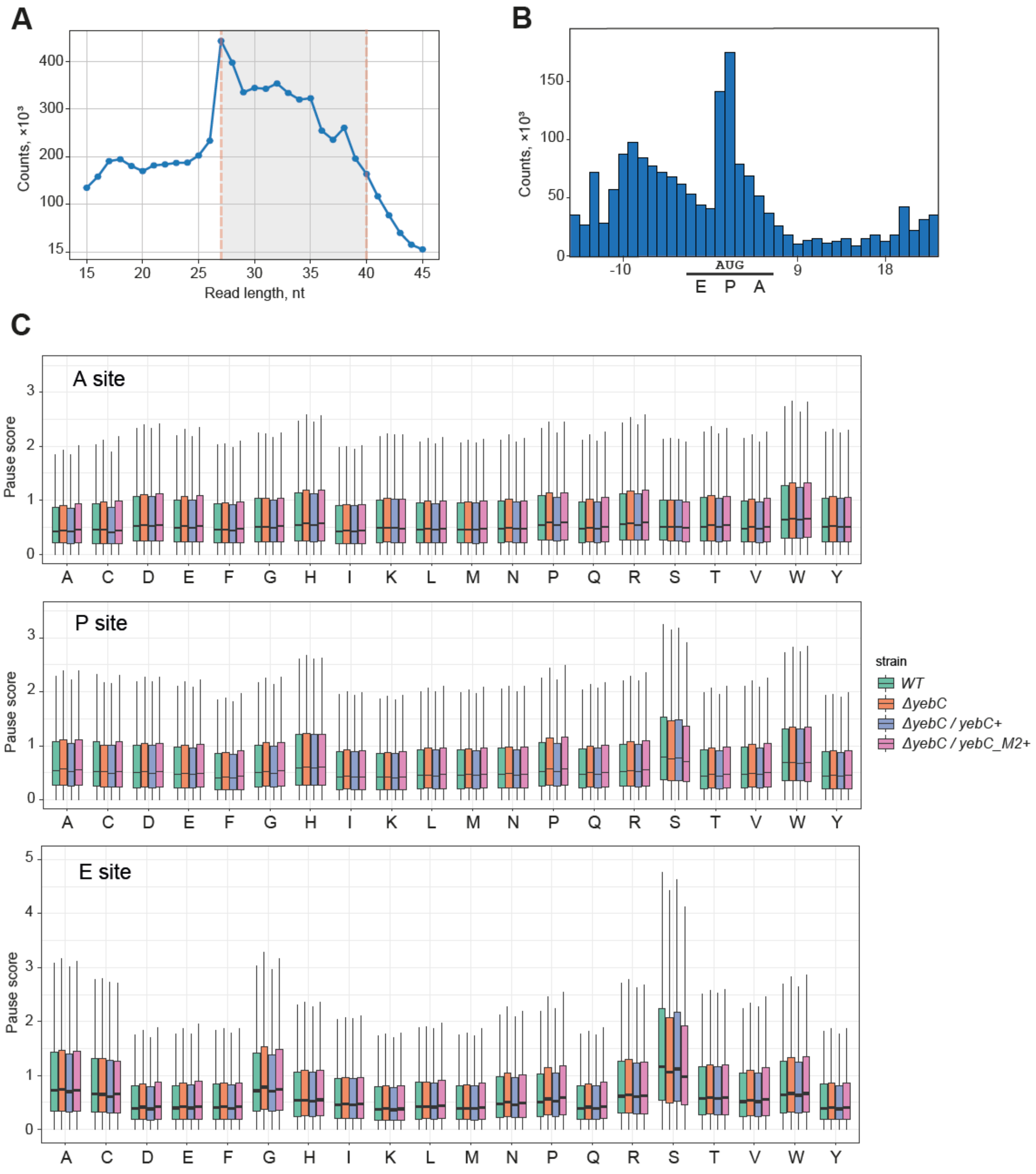


**Supplementary figure 8. Analysis of the ribosome profiling results.**

**(A)** Length distribution of mapped reads in the *WT* replicate 1 sample. The reads having lengths between 27 and 40 nucleotides were selected for further analysis.

**(B)** Assignment of the ribosome P site position near the start codon. The P site position for each mapped cDNA read was determined by shifting the 3′ end of the read by 15 nucleotides. The sum of the P site positions near the start codon in *WT* replicate 1 is presented. The increased number of P sites assigned to the start codon indicates the correct assignment of the ribosome position.

**(C)** Distribution of the pause scores for the codons in the A, P and E sites of the ribosome in the *WT* and *yebC* mutants. For each codon, the average pause score of three replicates was calculated in each sample. The distribution of the mean pause scores is represented by a boxplot, where the box represents the interquartile range and the median and the whiskers extend up to ×1.5 of the interquartile range.

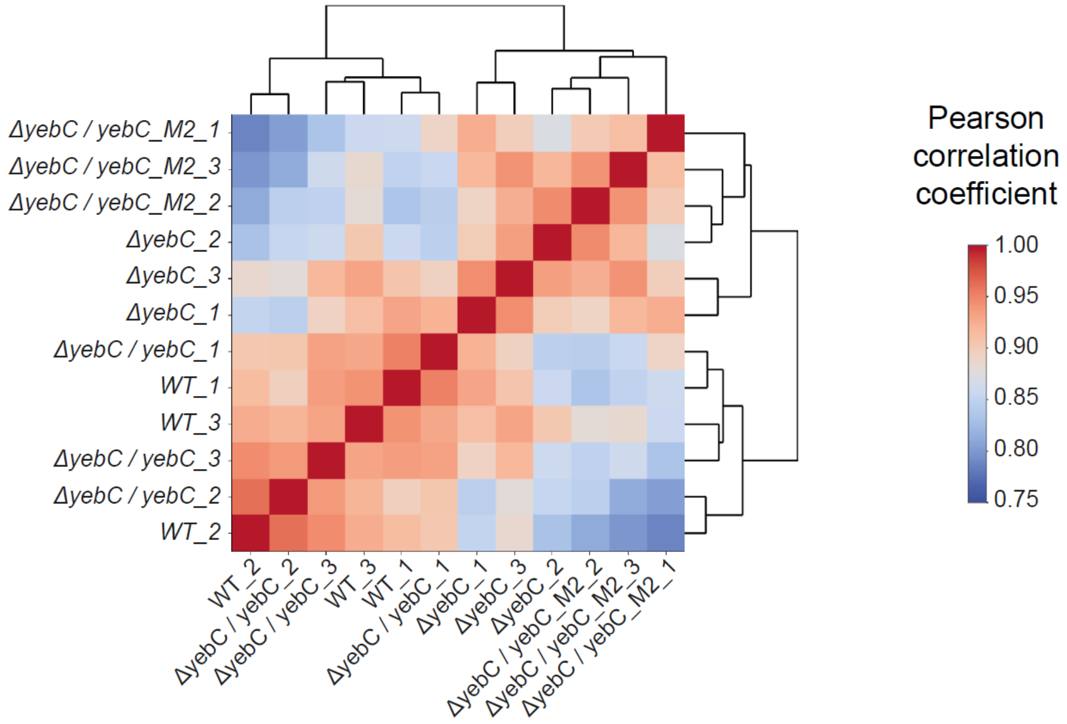


**Supplementary figure 9. Correlation between the ribosome profiling samples.**
The Pearson correlation coefficients were calculated between the pause scores of the ribosome profiling samples. The samples with high correlation coefficients are clustered together.

**pEC2817 (pSEVA141_3xFLAG-lox71-*erm*-lox66)**

lox71 lox66 ermAMB

```
<pSEVA141_backbone>GAATTCGAGCTCGGTACCCTACCGTTCGTATAGCATACATTATACGAAGTTATCCGTAGCGGTTTTCAAAATTTG
CAACCAGGAATGAATTACTATCCCTTTTATCAAGAAGCGCAAAAGAAAAACGAAATGATACACCAATCAGTGCAAAAAAAGATATAATGGGAGA
TAAGACGGTTCGTGTTCGTGCTGACTTGCACCATATCATAAAAATCGAAACAGCAAAGAATGGCGGAAACGTAAAAGAAGTTATGGAAATAAGA
CTTAGAAGCAAACTTAAGAGTGTGTTGATAGTGCAGTATCTTAAAATTTTGTATAATAGGAATTGAAGTTAAATTAGATGCTAAAAATTTGTAA
TTAAGAAGGAGTGATTACATGAACAAAAATATAAAATATTCTCAAAACTTTTTAACGAGTGAAAAAGTACTCAACCAAATAATAAAACAATTGA
ATTTAAAAGAAACCGATACCGTTTACGAAATTGGAACAGGTAAAGGGCATTTAACGACGAAACTGGCTAAAATAAGTAAACAGGTAACGTCTAT
TGAATTAGACAGTCATCTATTCAACTTATCGTCAGAAAAATTAAAACTGAATACTCGTGTCACTTTAATTCACCAAGATATTCTACAGTTTCAA
TTCCCTAACAAACAGAGGTATAAAATTGTTGGGAGTATTCCTTACCATTTAAGCACACAAATTATTAAAAAAGTGGTTTTTGAAAGCCATGCGT
CTGACATCTATCTGATTGTTGAAGAAGGATTCTACAAGCGTACCTTGGATATTCACCGAACACTAGGGTTGCTCTTGCACACTCAAGTCTCGAT
TCAGCAATTGCTTAAGCTGCCAGCGGAATGCTTTCATCCTAAACCAAAAGTAAACAGTGTCTTAATAAAACTTACCCGCCATACCACAGATGTT
CCAGATAAATATTGGAAGCTATATACGTACTTTGTTTCAAAATGGGTCAATCGAGAATATCGTCAACTGTTTACTAAAAATCAGTTTCATCAAG
CAATGAAACACGCCAAAGTAAACAATTTAAGTACCGTTACTTATGAGCAAGTATTGTCTATTTTTAATAGTTATCTATTATTTAACGGAGGAA
ATAAATAACTTCGTATAGCATACATTATACGAACGGTAGGGGATCCTCTAGAGTCGACCTGCAGGCATGCAAGCT<pSEVA141_backbone>
```

**pEC2818 (pSEVA141_3xFLAG-lox71-*erm*-lox66)**

C-terminal 3xFLAG lox71 lox66 ermAMB

```
<pSEVA141_backbone>GAATTCCCATGGGAGCTCGGTACCGACTACAAAGACCATGACGGTGATTATAAAGATCATGATATCGACTACAAA
GATGACGACGATAAATAGTAAGTACCCTACCGTTCGTATAGCATACATTATACGAAGTTATCCGTAGCGGTTTTCAAAATTTGCAACCAGGAAT
GAATTACTATCCCTTTTATCAAGAAGCGCAAAAGAAAAACGAAATGATACACCAATCAGTGCAAAAAAAGATATAATGGGAGATAAGACGGTTC
GTGTTCGTGCTGACTTGCACCATATCATAAAAATCGAAACAGCAAAGAATGGCGGAAACGTAAAAGAAGTTATGGAAATAAGACTTAGAAGCAA
ACTTAAGAGTGTGTTGATAGTGCAGTATCTTAAAATTTTGTATAATAGGAATTGAAGTTAAATTAGATGCTAAAAATTTGTAATTAAGAAGGAG
TGATTACATGAACAAAAATATAAAATATTCTCAAAACTTTTTAACGAGTGAAAAAGTACTCAACCAAATAATAAAACAATTGAATTTAAAAGAA
ACCGATACCGTTTACGAAATTGGAACAGGTAAAGGGCATTTAACGACGAAACTGGCTAAAATAAGTAAACAGGTAACGTCTATTGAATTAGACA
GTCATCTATTCAACTTATCGTCAGAAAAATTAAAACTGAATACTCGTGTCACTTTAATTCACCAAGATATTCTACAGTTTCAATTCCCTAACAA
ACAGAGGTATAAAATTGTTGGGAGTATTCCTTACCATTTAAGCACACAAATTATTAAAAAAGTGGTTTTTGAAAGCCATGCGTCTGACATCTAT
CTGATTGTTGAAGAAGGATTCTACAAGCGTACCTTGGATATTCACCGAACACTAGGGTTGCTCTTGCACACTCAAGTCTCGATTCAGCAATTGC
TTAAGCTGCCAGCGGAATGCTTTCATCCTAAACCAAAAGTAAACAGTGTCTTAATAAAACTTACCCGCCATACCACAGATGTTCCAGATAAATA
TTGGAAGCTATATACGTACTTTGTTTCAAAATGGGTCAATCGAGAATATCGTCAACTGTTTACTAAAAATCAGTTTCATCAAGCAATGAAACAC
GCCAAAGTAAACAATTTAAGTACCGTTACTTATGAGCAAGTATTGTCTATTTTTAATAGTTATCTATTATTTAACGGAGGAAATAAATAACTT
CGTATAGCATACATTATACGAACGGTAGGGGATCCTCTAGAGTCGACCTGCAGGCATGCAAGCTT<pSEVA141_backbone>
```

**pEC3160 (p7INTΔlacZα_pTet_3xFLAG-sfGFP-P0-mKate)**

tetR tetO 3xFLAG sfGFP mKate2

```
<p7INT_backbone>AAAAAAGGCCCACTTTTGTGGGCCTTTTTTTTAAGACCCACTTTCACATTTAAGTTGTTTTTCTAATCCGCAGATGAT
CAATTCAAGGCCGAATAAGAAGGCTGGCTCTGCACCTTGGTGATCAAATAATTCGATAGCTTGTCGTAATAATGGCGGCATACTATCAGTAGTA
GGTGTTTCCCTTTCTTCTTTAGCGACTTGATGCTCTTGATCTTCCAATACGCAACCTAAAGTAAAATGCCCCACAGCGCTGAGTGCATATAATG
CATTCTCTAGTGAAAAACCTTGTTGGCATAAAAAGGCTAATTGATTTTCGAGAGTTTCATACTGTTTTTCTGTAGGCCGTGTACCTAAATGTAC
TTTTGCTCCATCGCGATGACTTAGTAAAGCACATCTAAAACTTTTAGCGTTATTACGTAAAAAATCTTGCCAGCTTTCCCCTTCTAAAGGGCAA
AAGTGAGTATGGTGCCTATCTAACATCTCAATGGCTAAGGCGTCGAGCAAAGCCCGCTTATTTTTTACATGCCAATACAATGTAGGCTGCTCTA
CACCTAGCTTCTGGGCGAGTTTACGGGTTGTTAAACCTTCGATTCCGACCTCATTAAGCAGCTCTAATGCGCTGTTAATCACTTTACTTTTATC
TAATCTAGACATCATTAATTCCTCCTTTTTGTTGACACTCTATCATTGATAGAGTTATTTGTCAAACTAGTTTTTTATTTGGATCCCCTCGAGT
TCATGAAAAACTAAAAAAAATATTGACACTCTATCATTGATAGAGTATAATTAAAATAAGACTCTATCATTGATAGAGTCTTGATGGTACCGAG
CTCGAATAGATCTTCGAGTCTAGTTAAGGAGGTGATCTCATATGGACTACAAGGATCATGATGGTGATTATAAAGATCATGATATCGATTACAA
AGACGATGACGACAAGAGCAAAGGAGAAGAACTTTTCACTGGAGTTGTCCCAATTCTTGTTGAATTAGATGGTGATGTTAATGGGCACAAATTT
TCTGTCCGTGGAGAGGGTGAAGGTGATGCTACAAACGGAAAGCTTACCCTTAAATTTATTTGCACTACTGGAAAACTACCTGTTCCATGGCCAA
CACTTGTCACTACTCTCACTTATGGTGTTCAATGCTTTTCCCGTTATCCGGATCATATGAAACGGCATGACTTTTTCAAGAGTGCCATGCCCGA
AGGTTATGTACAGGAACGCACTATATCTTTCAAAGATGACGGGACTTACAAGACGCGTGCTGAAGTCAAGTTTGAAGGTGATACCCTTGTTAAT
CGTATCGAGTTAAAAGGTATTGATTTTAAAGAAGATGGAAACATTCTCGGACACAAACTTGAGTACAACTTTAACTCACACAATGTATACATCA
CGGCAGACAAACAAAAGAATGGAATCAAAGCTAACTTCAAAATTCGCCACAACGTTGAAGATGGATCCGTTCAACTAGCAGACCATTATCAACA
AAATACTCCAATTGGCGATGGCCCTGTCCTTTTACCAGACAACCATTACCTGTCGACACAATCTGTCCTTTCGAAAGATCCCAACGAAAAGCGT
GACCACATGGTCCTTCTTGAGTTTGTAACTGCTGCTGGGATTACACATGGCATGGATGAGCTCTACAAAGGATCTGTTTCAGAACTTATCAAAG
AAAACATGCACATGAAACTTTACATGGAAGGTACTGTTAACAACCACCACTTCAAATGTACTTCAGAAGGTGAAGGTAAACCATACGAAGGTAC
TCAAACTATGCGTATCAAAGCTGTTGAAGGTGGTCCACTTCCATTCGCTTTCGACATCCTTGCTACTTCATTCATGTACGGTTCAAAAACTTTC
ATCAACCACACTCAAGGTATCCCAGACTTCTTCAAACAATCATTCCCAGAAGGTTTCACTTGGGAACGTGTTACTACTTACGAAGACGGTGGTG
TTCTTACTGCTACTCAAGACACTTCACTTCAAGACGGTTGTCTTATCTACAACGTTAAAATCCGTGGTGTTAACTTCCCATCAAACGGTCCAGT
TATGCAAAAAAAAACTCTTGGTTGGGAAGCTTCAACTGAAACTCTTTACCCAGCTGACGGTGGTCTTGAAGGTCGTGCTGACATGGCTCTTAAA
CTTGTTGGTGGTGGTCACCTTATCTGTAACCTTAAAACTACTTACCGTTCAAAAAAACCAGCTAAAAACCTTAAAATGCCAGGTGTTTACTACG
TTGACCGTCGTCTTGAACGTATCAAAGAAGCTGACAAAGAAACTTACGTTGAACAACACGAAGTTGCTGTTGCTCGTTACTGTGACCTTCCATC
AAAACTTGGTCACCGTGCATAATAGTGACTGACGAAAAAAGGCCCACTTTTGTGGGCCTTTTTT<p7INT_backbone>
```

**Supplementary figure 10. Nucleotide sequences introduced to pSEVA141 and p7INT backbones.**

The partial sequences of the plasmids pSEVA141-lox71-erm-lox66, pSEVA141-3xFLAG-lox71-erm-lox66 and p7INTΔlacZα_pTet_3xFLAG-sfGFP-P0-mKate are presented. The coding sequences and other functional elements are indicated with colors.

### Supplementary tables

#### Supplementary table 1: Primers used in the study

| Code | Name | Sequence |
|---|---|---|
| **Introduction of 3x FLAG tag to the C-termini of *S. pyogenes* genes** | | |
| OLEC11928 | 3FLAG_yhaM_up_EcoRI_F | GATCAGAATTCCCTCAGGTTAATCAAATCACC |
| OLEC11929 | 3FLAG_yhaM_up_KpnI_R | GATCAGGTACCATAGTTTGGCTTGTAAAAAGACC |
| OLEC11930 | 3FLAG_yhaM_dw_BamHI_F | GATCAGGATCCTCAGTGTTTCTCGAGTAATAGTTC |
| OLEC11931 | 3FLAG_yhaM_dw_SalI_R | GATCAGTCGACCTACGCTTTGACAGAAACATC |
| OLEC11932 | 3FLAG_yhaM_seq_F | GGGCAATTAGAATACGGTAGTC |
| OLEC11933 | 3FLAG_yhaM_seq_R | TTGTTCTTATTAATATCATGCAAGAATTC |
| OLEC11934 | 3FLAG_gapN_up_EcoRI_F | GATCAGAATTCTTAAACCACCAACACAAGGCT |
| OLEC11935 | 3FLAG_gapN_up_KpnI_R | GATCAGGTACCCTGGATATCAAATACAACAGATTTAAC |
| OLEC11936 | 3FLAG_gapN_dw_BamHI_F | GATCAGGATCCTAAAAAAAATAAACAAAAAGTTAGGTTAAC |
| OLEC11937 | 3FLAG_gapN_dw_PstI_R | GATCACTGCAGGCCAAAATATCTAAAACTTGAGGA |
| OLEC11938 | 3FLAG_gapN_seq_F | GCATTGCTGAGCAATTAGAAG |
| OLEC11939 | 3FLAG_gapN_seq_R | AAACTATACGCTGACTATCTTCG |
| OLEC12918 | 3FLAG_yebC_up_NcoI_F | ACACACCATGGGTAAATGGGCAAATATTGTTGC |
| OLEC12919 | 3FLAG_yebC_up_KpnI_R | ACACAGGTACCAAAATCTGCTACATTATGATACAC |
| OLEC12920 | 3FLAG_yebC_dw_BamHI_F | ACACAGGATCCATTGACATAGAATAATAAAAAGAGTTG |
| OLEC12921 | 3FLAG_yebC_dw_PstI_R | ACACACTGCAGGCTCTGTTGAGGTGAGTTC |
| OLEC12922 | 3FLAG_yebC_seq_F | AAGTGCTATAATAAGGGAGTTAG |
| OLEC12923 | 3FLAG_ko_yebC_seq_R | TCCATCAGCAGTCAACTC |
| OLEC13855 | 3FLAG_phoH_up_NcoI_F | ACACACCATGGGCTTTATCTATGGCAGAATCTC |
| OLEC13856 | 3FLAG_phoH_up_KpnI_R | ACACAGGTACCGTGCTCTTGACCGATCAC |
| OLEC13857 | 3FLAG_phoH_dw_XbaI_F | ACACATCTAGAGACTTTTACGATGATGTTTAATATGG |
| OLEC13858 | 3FLAG_phoH_dw_PaeI_R | ACACAGCATGCCCCTTATCCTTTCAATCAGC |
| OLEC13941 | 3FLAG_phoH_seq_F | GTGATGATGAAGAAGCTG |
| OLEC13942 | 3FLAG_phoH_seq_R | AATCATCTCGATATACATAAGG |
| OLEC13859 | 3FLAG_thuC_up_NcoI_F | ACACACCATGGTTTCTTTTGATGTGCCTGAC |
| OLEC13860 | 3FLAG_thuC_up_KpnI_R | ACACAGGTACCTGAGTTTTTCTTAACATTTTCTATAATCG |
| OLEC13861 | 3FLAG_thuC_dw_BamHI_F | ACACAGGATCCTGTGGGTGAATTTGGGG |
| OLEC13862 | 3FLAG_thuC_dw_SalI_R | ACACAGTCGACTCGATATCATTGCTGGATAG |
| OLEC13943 | 3FLAG_thuC_seq_F | AATTTAATCTTGTCAACTCAAC |
| OLEC13944 | 3FLAG_thuC_seq_R | CATTCTACCTCTGACAGC |
| OLEC12950 | 3FLAG_yjbK_up_NcoI_F | ACACACCATGGGTCTCCTTTCTTGTCAAAGC |
| OLEC12951 | 3FLAG_yjbK_up_KpnI_R | ACACAGGTACCTTTATCGTTGAACTTTTTTAAGGTA |
| OLEC12952 | 3FLAG_yjbK_dw_BamHI_F | ACACAGGATCCTGGCTAAAAGCGACAGAAAAAC |
| OLEC12953 | 3FLAG_yjbK_dw_PstI_R | ACACACTGCAGTTTAGCAGCTTCTGCAAAGG |
| OLEC12954 | 3FLAG_yjbK_seq_F | CGATATTGCTTGCGAATACC |
| OLEC12955 | 3FLAG_yjbK_seq_R | CTCCACCAGCAAATAAGC |

| Code | Name | Sequence |
|---|---|---|
| OLEC12966 | 3FLAG_ygaC_up_NcoI_F | ACACACCATGGGCGACGACATTAATTGCAAG |
| OLEC12967 | 3FLAG_ygaC_up_KpnI_R | ACACAGGTACCACGATTCTTCAGTTCAAGATAAC |
| OLEC12968 | 3FLAG_ygaC_dw_BamHI_F | ACACAGGATCCGAGTTGTCCAGCACTCC |
| OLEC12969 | 3FLAG_ygaC_dw_PstI_R | ACACACTGCAGTTTGTCTGTTTTACGAACGTG |
| OLEC12970 | 3FLAG_ygaC_seq_F | GTCGCAAATATGATGGTTATACC |
| OLEC12971 | 3FLAG_ygaC_seq_R | CTTGGAAAAGTTTATGCAGAC |
| **Knock-out of *yebC* in *S. pyogenes*** | | |
| OLEC13193 | ko_yebC_up_EcoRI_F | ACACAGAATTCAAACGCTATACAAAAGCTCG |
| OLEC13194 | ko_yebC_up_KpnI_R | ACACAGGTACCTTTTGTCTCCTTTTAATAGTATTTTATTG |
| OLEC12920 | ko_yebC_dw_BamHI_F | ACACAGGATCCATTGACATAGAATAATAAAAAGAGTTG |
| OLEC13195 | ko_yebC_dw_SalI_R | ACACAGTCGACACTCAAATTGGTTAACATTTGAG |
| OLEC13196 | ko_yebC_seq_F | AATGATGAATTGGCAAGTCG |
| OLEC12923 | 3FLAG_ko_yebC_seq_R | TCCATCAGCAGTCAACTC |
| **Complementation of *yebC* in *S. pyogenes*** | | |
| OLEC13350 | comp_p7INT_BamHI_F | ACACAGGATCCCATTAATGAATCGGCCAAC |
| OLEC13351 | comp_p7INT_XbaI_R | ACACATCTAGACGTAATAGCGAAGAGGC |
| OLEC13346 | comp_yebC_BamHI_F | ACACAGGATCCGCTGTTGTCACTCAAGC |
| OLEC13347 | comp_yebC_XbaI_R | ACACATCTAGACAACGACTATGTTACCTGG |
| OLEC13348 | comp_yebC_3FLAG_F | TCATGATATCGACTACAAAGATGACGACGATAAATAGTAAATTGACATAGA ATAATAAAAAG |
| OLEC13349 | comp_yebC_3FLAG_R | TCTTTATAATCACCGTCATGGTCTTTGTAGTCGGTACCAAAATCTGCTACA TTATGATA |
| **Site-directed mutagenesis of *yebC* in *S. pyogenes*** | | |
| OLEC14266 | SDM_yebC_m1_F | TGCCGCATTTGGTGTTGAAATTTATGTG |
| OLEC14267 | SDM_yebC_m1_R | TAGACAGCTGATGTTGCTCCATCTTTAG |
| OLEC14268 | SDM_yebC_m3_F | GCTGCGATTGATGCAGCTAAAGGAAACACAGATGAAAC |
| OLEC14269 | SDM_yebC_m3_R | ATCAATAACTGCAGCTGGCACTTGTGCTTGCTT |
| OLEC14270 | SDM_yebC_m4_F | GCAACGGCTTACGGTGCTAACGGTGGCAATATGGGA |
| OLEC14271 | SDM_yebC_m4_R | TACATTTGCCGCTGTAGCGTTAACATTTGATGTCAAAGTATCC |
| OLEC14388 | SDM_yebC_m5_F | TGCACAATTACTTGCTGCGGATGTAGACGTAGATG |
| OLEC14389 | SDM_yebC_m5_R | AAGACAGCTGCAGCAGCATCACCAGCAAAAACGATG |
| OLEC14392 | SDM_yebC_m6_F | TGCAACTTTTGCAAAGCTTATTGATGCACTTG |
| OLEC14393 | SDM_yebC_m6_R | AGGTCAGCACCTGCCAAAACTACTTCTGATTGAG |
| OLEC14272 | SDM_yebC_Y84A_F | AGAGGGACGCGCTGAAGGTTTTG |
| OLEC14273 | SDM_yebC_Y84A_Y84F_R | ACGAAAGTTTCATCTGTG |
| OLEC14387 | SDM_yebC_Y84F_F | AGAGGGACGCTTTGAAGGTTTTG |
| OLEC14274 | SDM_yebC_E85A_F | GGGACGCTATGCTGGTTTTGGTC |
| OLEC14275 | SDM_yebC_E85A_R | TCTACGAAAGTTTCATCTG |
| **iCLIP2 for YebC in *S. pyogenes*** | | |
| OLEC14114 | RToligo | GGATCCTGAACCGCT |
| OLEC14115 | L3-App | /rApp/AGATCGGAAGAGCGGTTCAG/ddC/ |

| Code | Name | Sequence |
|---|---|---|
| OLEC14167 | L07clip2.0 | /5Phos/NNNNCAGATCNNNNNAGATCGGAAGAGCGTCGTG/3ddC/ |
| OLEC14168 | L08clip2.0 | /5Phos/NNNNACTTGANNNNNAGATCGGAAGAGCGTCGTG/3ddC/ |
| OLEC14169 | L09clip2.0 | /5Phos/NNNNGATCAGNNNNNAGATCGGAAGAGCGTCGTG/3ddC/ |
| OLEC14170 | L10clip2.0 | /5Phos/NNNNTAGCTTNNNNNAGATCGGAAGAGCGTCGTG/3ddC/ |
| OLEC14171 | L11clip2.0 | /5Phos/NNNNATGAGCNNNNNAGATCGGAAGAGCGTCGTG/3ddC/ |
| OLEC14172 | L12clip2.0 | /5Phos/NNNNCTTGTANNNNNAGATCGGAAGAGCGTCGTG/3ddC/ |
| OLEC14120 | P5Solexa_s | ACACGACGCTCTTCCGATCT |
| OLEC14121 | P3Solexa_s | CTGAACCGCTCTTCCGATCT |
| OLEC14122 | P5Solexa | AATGATACGGCGACCACCGAGATCTACACTCTTTCCCTACACGACGCTCT TCCGATCT |
| OLEC14123 | P3Solexa | CAAGCAGAAGACGGCATACGAGATCGGTCTCGGCATTCCTGCTGAACCG CTCTTCCGATCT |
| OLEC14202 | Seq_1 | ACACTCTTTCCCTACACGACGCTCTTCCGATCT |
| OLEC14203 | Seq_2 | CGGTCTCGGCATTCCTGCTGAACCGCTCTTCCGATCT |
| **Ribosome profiling** | | |
| OLEC12113 | ribseq_15_control (RNA) | AUGUACACGGAGUCG |
| OLEC12114 | ribseq_45_control (RNA) | AUGUACACGGAGUCGACCCGCAACGCGAUGUACACGGAGUCGACC |
| OLEC12115 | ribseq_adapter_with_UMI | /rApp/NNNNNATCGTAGATCGGAAGAGCACACGTCTGAA/3ddC/ |
| OLEC12116 | ribseq_RT_primer | /5Phos/NNAGATCGGAAGAGCGTCGTGTAGGGAAAGAG/iSp18/GTGACTG GAGTTCAGACGTGTGCTC |
| OLEC12244 | ribseq_libamp_fw | AATGATACGGCGACCACCGAGATCTACACTCTTTCCCTACACGACGCTC |
| OLEC12245 | ribseq_libamp_rev_ind_1 | CAAGCAGAAGACGGCATACGAGATCGTGATGTGACTGGAGTTCAGACGT GTG |
| OLEC12246 | ribseq_libamp_rev_ind_2 | CAAGCAGAAGACGGCATACGAGATACATCGGTGACTGGAGTTCAGACGT GTG |
| OLEC12247 | ribseq_libamp_rev_ind_3 | CAAGCAGAAGACGGCATACGAGATGCCTAAGTGACTGGAGTTCAGACGT GTG |
| OLEC14922 | ribseq_libamp_rev_ind_4 | CAAGCAGAAGACGGCATACGAGATTGGTCAGTGACTGGAGTTCAGACGT GTG |
| OLEC14923 | ribseq_libamp_rev_ind_5 | CAAGCAGAAGACGGCATACGAGATCACTGTGTGACTGGAGTTCAGACGT GTG |
| OLEC14924 | ribseq_libamp_rev_ind_6 | CAAGCAGAAGACGGCATACGAGATATTGGCGTGACTGGAGTTCAGACGT GTG |
| OLEC14925 | ribseq_libamp_rev_ind_7 | CAAGCAGAAGACGGCATACGAGATGATCTGGTGACTGGAGTTCAGACGT GTG |
| OLEC14926 | ribseq_libamp_rev_ind_8 | CAAGCAGAAGACGGCATACGAGATTCAAGTGTGACTGGAGTTCAGACGT GTG |
| OLEC14927 | ribseq_libamp_rev_ind_9 | CAAGCAGAAGACGGCATACGAGATCTGATCGTGACTGGAGTTCAGACGT GTG |
| OLEC14928 | ribseq_libamp_rev_ind_10 | CAAGCAGAAGACGGCATACGAGATAAGCTAGTGACTGGAGTTCAGACGT GTG |
| OLEC14929 | ribseq_libamp_rev_ind_11 | CAAGCAGAAGACGGCATACGAGATGTAGCCGTGACTGGAGTTCAGACGT GTG |
| OLEC14930 | ribseq_libamp_rev_ind_12 | CAAGCAGAAGACGGCATACGAGATTACAAGGTGACTGGAGTTCAGACGT GTG |
| **Ribosome stalling reporter in *S. pyogenes*** | | |
| OLEC15067 | P5_P3_R | TGGAGGTGGTCCTTTGTAGAGCTCATCCATG |
| OLEC15068 | P5_F | CCTCCATCTGTTTCAGAACTTATCAAAGAAAAC |
| OLEC15120 | link_F | TCTGTTTCAGAACTTATCAAAGAAAACATGC |
| OLEC15122 | stop_R | TTATCCTTTGTAGAGCTCATCCATGCCATGTG |
| OLEC15140 | PPG_R | TCCAGGTGGTCCTTTGTAGAGCTC |
| OLEC15141 | PIP_R | TGGAATTGGTCCTTTGTAGAGCTCATCC |
| **Ribosome stalling reporter *in vitro*** | | |

| Code | Name | Sequence |
|---|---|---|
| OLEC14889 | HiFi_p21_F | CATATGTATATCTCCTTCTTAAAGTTAAAC |
| OLEC14890 | HiFi_p21_R | CTCGAGCACCACCACCAC |
| OLEC14953 | HiFi_b1983_F | TTAAGAAGGAGATATACATATGGGACGTAAATGGGCC |
| OLEC14954 | HiFi_b1983_R | AGTGGTGGTGGTGGTGGTGCTCGAGGAGATTTGCGACGTTATGATAAAC |
| OLEC15057 | HiFi_b1864_F | TTAAGAAGGAGATATACATATGGCAGGTCATAGTAAATGGGC |
| OLEC15058 | HiFi_b1864_R | AGTGGTGGTGGTGGTGGTGCTCGAGGAGAGTCGCTGCGACC |
| OLEC15015 | b1983_HA_F | ATCGTATGGGTAGGTACCGAGATTTGCGACGTTATGATAAAC |
| OLEC15016 | b1983_b1864_HA_R | GTTCCAGATTACGCTCATCACCACCACCACCACTGAG |
| OLEC15093 | b1864_HA_F | ATCGTATGGGTAGGTACCGAGAGTCGCTGCGACCTC |
| OLEC15063 | b1983_m2_1_F | GCTGCAATTGATGCAGCCAAAGGCGGCG |
| OLEC15064 | b1983_m2_1_R | ATCAATAACAGCTGCTGGAACTTGTGCCTGC |
| OLEC15065 | b1983_m2_2_F | ACAATTTTCAATGCAAAAGGCGGCAATATCGG |
| OLEC15066 | b1983_m2_2_R | TGCAACGTTAGCAATCGTAGCGTTAACATTTGAAGTC |
| OLEC15073 | b1983_Y84A_F | GCAGAAGGCTTTGGTCCTAATGGC |
| OLEC15074 | b1983_Y84A_R | ACGTCCCTGCACGAAC |
| OLEC15051 | folA_FLAG_F | ATCAGTCTGATTGCGGC |
| OLEC14962 | folA_FLAG_R | CTTGTCGTCATCGTCTTTGTAGTCCATATGTATATCTCCTTCTTAAAGTTAAAC |
| OLEC15076 | folA_P0_F | GCGGGACGCAAAAATATTATCCTCAGC |
| OLEC15165 | folA_P0_R | CAAGGCACGACCGATTGATTCCCAG |
| OLEC15054 | folA_P5_F | TGGTGGCGGACGACCGATTGATTC |
| OLEC15055 | folA_P5_R | CCGCCAGGACGCAAAAATATTATCCTCAG |
| OLEC14961 | folA_stop_F | TAATGAGGATCCCGGGAATTC |
| OLEC15052 | folA_stop_R | CAACGGACGACCGATTGATTC |

#### Supplementary table 2: Plasmids used in the study

| Code | Description |
|---|---|
| **Introduction of 3x FLAG tag to the C-termini of *S. pyogenes* genes** | |
| pEC2818 | pSEVA141-3xFLAG-lox71-erm-lox66 |
| pEC2819 | pSEVA141-yhaM-3xFLAG-lox71-erm-lox66 |
| pEC2820 | pSEVA141-gapN-3xFLAG-lox71-erm-lox66 |
| pEC2969 | pSEVA141-yebC-3xFLAG-lox71-erm-lox66 |
| pEC2972 | pSEVA141-yjbK-3xFLAG-lox71-erm-lox66 |
| pEC2974 | pSEVA141-ygaC-3xFLAG-lox71-erm-lox66 |
| pEC3034 | pSEVA141-phoH-3xFLAG-lox71-erm-lox66 |
| pEC3035 | pSEVA141-thuC-3xFLAG-lox71-erm-lox66 |
| pEC455 | repDEG-pAMbeta1-pJH1-aphIII-bgaB-colE1-PgyrA-Cre |
| ***yebC* mutants in *S. pyogenes*** | |
| pEC2817 | pSEVA141-lox71-erm-lox66 |
| pEC2976 | pSEVA141-yebC_lox71-erm-lox66 |
| pEC2988 | p7INTΔlacZα_Spy0316-3xFLAG |
| pEC3071 | p7INTΔlacZα_Spy0316(K21A, K25A)-3xFLAG |
| pEC3085 | p7INTΔlacZα_Spy0316(K61A, H62A, K66A, K70A, R105A, R111A, K116A)-3xFLAG |
| pEC3072 | p7INTΔlacZα_Spy0316(K61A, H62A, K66A, K70A)-3xFLAG |
| pEC3073 | p7INTΔlacZα_Spy0316(R105A, R111A, K116A)-3xFLAG |
| pEC3087 | p7INTΔlacZα_Spy0316(D143A, D145A, S146A, E149A, E153A)-3xFLAG |
| pEC3089 | p7INTΔlacZα_Spy0316(E208A, D210A, E213A, E216A)-3xFLAG |
| pEC3074 | p7INTΔlacZα_Spy0316(Y84A)-3xFLAG |
| pEC3086 | p7INTΔlacZα_Spy0316(Y84F)-3xFLAG |
| pEC3075 | p7INTΔlacZα_Spy0316(E85A)-3xFLAG |
| **Ribosome stalling reporter in *S. pyogenes*** | |
| pEC3155 | p7INTΔlacZα_pTet_3xFLAG-sfGFP-P5-mKate |
| pEC3156 | p7INTΔlacZα_pTet_3xFLAG-sfGFP-P3-mKate |
| pEC3157 | p7INTΔlacZα_pTet_3xFLAG-sfGFP-PPG-mKate |
| pEC3158 | p7INTΔlacZα_pTet_3xFLAG-sfGFP-PIP-mKate |
| pEC3160 | p7INTΔlacZα_pTet_3xFLAG-sfGFP-P0-mKate |
| pEC3161 | p7INTΔlacZα_pTet_3xFLAG-sfGFP-stop-mKate |
| **Ribosome stalling reporter *in vitro*** | |
| pEC3162 | pET21-a(+)_b1983:HA tag |
| pEC3163 | pET21-a(+)_b1983(K61A, H62A, K66A, K70A, R105A, R111A, K116A):HA tag |
| pEC3164 | pET21-a(+)_b1983(Y84A):HA tag |
| pEC3165 | pET21-a(+)_b1864:HA tag |

#### Supplementary table 3: Strains of *S. pyogenes* used in the study

| Code | Strain | Genotype |
|---|---|---|
| EC2224 | *WT* | SF370 (M1 serotype) |
| EC3638 | *yhaM:3xFLAG* | SPy_0267::3xFLAG-lox72 |
| EC3639 | *gapN:3xFLAG* | SPy_1371::3xFLAG-lox72 |
| EC3640 | *yebC:3xFLAG* | SPy_0316::3xFLAG-lox72 |
| EC3646 | *phoH:3xFLAG* | SPy_0471::3xFLAG-lox72 |
| EC3647 | *thuC:3xFLAG* | SPy_0539::3xFLAG-lox72 |
| EC3707 | *yjbK:3xFLAG* | SPy_1124::3xFLAG-lox72 |
| EC3641 | *ygaC:3xFLAG* | SPy_1608::3xFLAG-lox72 |
| EC3615 | *ΔyebC* | ΔSPy_0316::lox72 |
| EC3619 | *Δyebc / yebC+* | ΔSPy_0316::lox72 SPy_S01::p7INTΔlacZα_SPy_0316::3xFLAG |
| EC3690 | *Δyebc / yebC_M1+* | ΔSPy_0316::lox72 SPy_S01::p7INTΔlacZα_SPy_0316(K21A, K25A):3xFLAG |
| EC3691 | *Δyebc / yebC_M3+* | ΔSPy_0316::lox72 SPy_S01::p7INTΔlacZα_SPy_0316(K61A, H62A, K66A, K70A):3xFLAG |
| EC3692 | *Δyebc / yebC_M4+* | ΔSPy_0316::lox72 SPy_S01::p7INTΔlacZα_SPy_0316(R105A, R111A, K116A):3xFLAG |
| EC3693 | *Δyebc / yebC_Y84A+* | ΔSPy_0316::lox72 SPy_S01::p7INTΔlacZα_SPy_0316(Y84A):3xFLAG |
| EC3694 | *Δyebc / yebC_E85A+* | ΔSPy_0316::lox72 SPy_S01::p7INTΔlacZα_SPy_0316(E85A):3xFLAG |
| EC3714 | *Δyebc / yebC_M2+* | ΔSPy_0316::lox72 SPy_S01::p7INTΔlacZα_SPy_0316(K61A, H62A, K66A, K70A, R105A, R111A, K116A):3xFLAG |
| EC3715 | *Δyebc / yebC_Y84F+* | ΔSPy_0316::lox72 SPy_S01::p7INTΔlacZα_SPy_0316(Y84F):3xFLAG |
| EC3716 | *Δyebc / yebC_M5+* | ΔSPy_0316::lox72 SPy_S01::p7INTΔlacZα_SPy_0316(D143A, D145A, S146A, E149A, E153A):3xFLAG |
| EC3718 | *Δyebc / yebC_M6+* | ΔSPy_0316::lox72 SPy_S01::p7INTΔlacZα_SPy_0316(E208A, D210A, E213A, E216A):3xFLAG |
| EC3788 | *WT sfGFP-P5-mKate* | Spy_s01::p7INTΔlacZα_pTet_3xFLAG-sfGFP-P5-mKate |
| EC3789 | *WT sfGFP-P3-mKate* | Spy_s01::p7INTΔlacZα_pTet_3xFLAG-sfGFP-P3-mKate |
| EC3790 | *WT sfGFP-PPG-mKate* | Spy_s01::p7INTΔlacZα_pTet_3xFLAG-sfGFP-PPG-mKate |
| EC3791 | *WT sfGFP-PIP-mKate* | Spy_s01::p7INTΔlacZα_pTet_3xFLAG-sfGFP-PIP-mKate |
| EC3794 | *ΔyebC sfGFP-P5-mKate* | ΔSPy_0316 Spy_s01::p7INTΔlacZα_pTet_3xFLAG-sfGFP-P5-mKate |
| EC3795 | *ΔyebC sfGFP-P3-mKate* | ΔSPy_0316 Spy_s01::p7INTΔlacZα_pTet_3xFLAG-sfGFP-P3-mKate |
| EC3796 | *ΔyebC sfGFP-PPG-mKate* | ΔSPy_0316 Spy_s01::p7INTΔlacZα_pTet_3xFLAG-sfGFP-PPG-mKate |
| EC3797 | *ΔyebC sfGFP-PIP-mKate* | ΔSPy_0316 Spy_s01::p7INTΔlacZα_pTet_3xFLAG-sfGFP-PIP-mKate |
| EC3799 | *ΔyebC sfGFP-P0-mKate* | ΔSPy_0316 Spy_s01::p7INTΔlacZα_pTet_3xFLAG-sfGFP-P0-mKate |
| EC3800 | *ΔyebC sfGFP-stop-mKate* | ΔSPy_0316 Spy_s01::p7INTΔlacZα_pTet_3xFLAG-sfGFP-stop-mKate |

### Supplementary methods

#### Mutagenesis of *S. pyogenes*

***Introduction of 3x FLAG to the C-termini of S. pyogenes genes***

The introduction of 3x FLAG to the C-termini of *S. pyogenes* genes was performed using the Cre-Lox recombination system. The *S. pyogenes* suicidal plasmid pSEVA141_3xFLAG-lox71-*erm*-lox66 (Supplementary fig. 10) encodes the 3xFLAG tag sequence and the lox71-*erm*-lox66 cassette in the pSEVA141 backbone. The 3xFLAG sequence is N-terminally flanked by the KpnI restriction site allowing in-frame cloning of coding sequences and terminates with a stop codon. For each gene the sequences upstream and downstream of their stop codons were amplified from *S. pyogenes* genomic DNA using primers listed in Supplementary table 1. The primers introduced the cutting sites for restriction enzymes. To generate the plasmids for the 3xFLAG introduction (Supplementary table 2), the upstream shoulders were cloned in-frame with the 3x FLAG, and the downstream shoulders were cloned downstream of the lox71-*erm*-lox66 cassette using the restriction enzymes.

The competent *S. pyogenes* cells for transformation with the suicidal vectors were prepared. The cells were grown in THY supplemented with 250 mM sucrose (Calbiochem) and 40 mM L-threonine (Sigma-Aldrich) until the midlogarithmic growth phase. The cells were washed with 0.5 M sucrose and resuspended in 0.5 M sucrose and 20% glycerol. 15 µg of the plasmids for 3xFLAG introduction were digested with Cfr42I and introduced by electroporation in a 0.1 cm electrode gap cuvette with a 1.8 kV, 400Ω and 25 µF pulse. After 2 h of growth in 5 mL of THY, the cells were plated on TSA plates with 2.5 µg/mL erythromycin. After two days the individual colonies were restreaked to the fresh erythromycin plates and incubated for an additional day. The regrown colonies were inoculated into fresh THY and grown over day, the clones were collected and the glycerol stocks were prepared.

For excision of the lox71-*erm*-lox66 cassette, the clones were transformed with the replicative pEC455 plasmid (Supplementary table 2) encoding the Cre recombinase. The *S. pyogenes* cells were grown in THY until the mid-logarithmic growth phase. The cells were washed with ice-cold mQ, resuspended in 20% glycerol and electroporated with 0.5 µg of the pEC455 plasmid. After 2 h of growth in 5 mL of THY the cells were plated on TSA plates with 250 µg/mL kanamycin. After two days, the individual colonies were inoculated into 2 mL of THY and grown over day to remove the pEC455 plasmid. The cultures were streaked on TSA plates without antibiotics and allowed to grow overnight. The colonies were inoculated into fresh THY and grown over day, the clones were collected and the glycerol stocks were prepared. The loss of the *erm* resistance cassette and pEC455 was confirmed by streaking the cultures on erythromycin and kanamycin plates. The introduction of C-terminal 3x FLAG followed by the lox72 site was confirmed by isolation of genomic DNA with NucleoSpin Microbial DNA kit (Macherey-Nagel), amplification of the corresponding genomic regions with primers listed in Supplementary table 1 and Sanger sequencing of the PCR fragments with the same primers.

***Deletion and complementation of yebC in S. pyogenes***

The deletion of the *yebC* gene in *S. pyogenes* was carried out using the Cre-Lox recombination system and the protocol is similar to the introduction of 3x FLAG to the C-termini of genes. The *S. pyogenes* suicidal plasmid pSEVA141_lox71-*erm*-lox66 (Supplementary fig. 10) bears the lox71-*erm*-lox66 cassette in the pSEVA141 backbone. The sequences upstream and downstream of *yebC* (*SPy_0316*) were amplified from *S. pyogenes* genomic DNA using primers listed in Supplementary table 1. The primers introduced the cutting sites for restriction enzymes. Using the restriction enzymes, the fragments were cloned upstream and downstream of the lox71-*erm*-lox66 cassette to generate the plasmid for *yebC* deletion (Supplementary table 2).

The WT and mutant versions of *yebC* were introduced into the *S. pyogenes ΔyebC* strain with the modified p7INT integrative vector. We used a version of p7INT with deleted *lacZ* and its promoter to avoid the non-specific transcription and decrease the plasmid size. The p7INT portion without the *lacZ* promotor and coding sequence, and *yebC* locus including the promotor and terminator were amplified with the primers listed in Supplementary table 1. The primers introduced the cutting sites for restriction enzymes that were used for cloning. Next, the 3x FLAG sequence was introduced to the C-terminus of *yebC* by the whole-plasmid PCR and circularization with KLD Enzyme Mix (New England Biolabs). The mutations to the *yebC* sequence were introduced by the whole-plasmid PCR with respective primers followed by circularization with KLD Enzyme Mix. The M2 mutation is a combination of M3 and M4 and was created by two rounds of PCR mutagenesis.

The substitution of the *yebC* gene in *S. pyogenes* genome with the lox71-*erm*-lox66 cassette and excision of the cassette were performed according to the same protocol as for the introduction of 3x FLAG to the C-termini of the genes. The deletion was confirmed by isolation of genomic DNA using the NucleoSpin Microbial DNA kit (Macherey-Nagel), amplification of *yebC* genomic locus with primers listed in Supplementary table 1 and Sanger sequencing of the PCR fragments with the same primers.

The competent cells of the *ΔyebC* strain were prepared in the same way as for the transformation with the pEC455 plasmid. The cells were transformed with 0.5 µg of p7INT derivatives. After 2 h of growth in 5 mL of THY the cells were plated on TSA plates with 2.5 µg/mL erythromycin. The colonies were restreaked onto the fresh erythromycin plates and incubated for an additional day. The regrown colonies were inoculated into fresh THY and grown over day, the clones were collected and the glycerol stocks were prepared.

***Generation of S. pyogenes reporter strains***

The plasmid p7INTΔlacZα_pTet_3xFLAG-sfGFP-P0-mKate encodes the fusion protein 3x FLAG-sfGFP-mKate2 under the control of the pTet promoter in the p7INT backbone (Supplementary fig. 10). The sfGFP and mKate amino acid sequences are connected via a Gly-Ser (P0) linker. To change the linker sequence, the whole-plasmid PCR was performed with the following primers: (P5) P5_F and P5_P3_R; (P3) link_F and P5_P3_R; (PPG) link_F and PPG_R; (PIP) link_F and PIP_R; (stop) link_F and stop_R (Supplementary table 1). The products were circularized with the KLD enzyme mix (New England Biolabs). The resulting vectors (Supplementary table 2) were transformed into the wild-type and *ΔyebC* strains using the same protocol as for the complementation with the *yebC* protein.

#### Mass spectrometry

***Mass spectrometry of OOPS samples***

The samples were analysed on an Orbitrap Fusion Lumos (Thermo Scientific) that was coupled to a 3000 RSLCnano UPLC (Thermo Scientific). Samples were loaded on a PepMap Trap cartridge (300 µm i.d. x 5 mm, C18, Thermo Scientific) with 2% acetonitrile, 0.1% TFA at a flow rate of 20 µL/min. Peptides were separated over a 50 cm analytical column (PicoFrit, 360 µm O.D., 75 µm I.D., 10 µm tip opening, non-coated, New Objective) that was packed in-house with Poroshell 120 EC-C18, 2.7 µm (Agilent). Solvent A consists of 0.1% formic acid in water. Elution was carried out at a constant flow rate of 250 nL/min using a 180-min method: 8-33% solvent B (0.1% formic acid in 80% acetonitrile) within 120 min, 33-48% solvent B within 25 min, 48-98% buffer B within 1 min, followed by column washing and equilibration. The Orbitrap Fusion Lumos mass spectrometer was equipped with a FAIMS Pro device, which was operated at standard resolution using three alternating CVs of -40V, -60V and -80V (cycle time for each was set to 1 second). Data acquisition was carried out using data-dependent acquisition in positive ion mode. MS survey scans were acquired from 375-1500 m/z in profile mode at a resolution of 240,000. AGC target was set to $4e^5$ charges, allowing a maximum injection time of 50 ms. Peptides with charge states 2-6 were subjected to CID fragmentation (fixed CE = 35%, AGC = $1e^4$) and analysed in the linear ion trap at a resolution of 125,000 Da/second. The isolation window was set to 1.6 m/z.

***Mass spectrometry of RBS-ID samples***

The samples were analysed on an Orbitrap Exploris 480 (Thermo Scientific) and Orbitrap Fusion Lumos, which were both coupled to a 3000 RSLCnano UPLC (Thermo Scientific). Samples were loaded on a PepMap Trap cartridge (300 µm i.d. x 5 mm, C18, Thermo Scientific) with 2% acetonitrile,

0.1% TFA at a flow rate of 20 µL/min. Peptides were separated over a 50 cm analytical column (PicoFrit, 360 µm O.D., 75 µm I.D., 10 µm tip opening, non-coated, New Objective) that was packed in-house with Poroshell 120 EC-C18, 2.7 µm (Agilent). Solvent A consists of 0.1% formic acid in water. Settings for Exploris: Elution was carried out at a constant flow rate of 250 nL/min within 90 min. Initially, a two-step linear gradient was applied: 3-30% solvent B (0.1% formic acid in 80% acetonitrile) within 74 min, 30-45% solvent B within 14 min, followed by column washing and equilibration. Data acquisition was carried out using data-dependent acquisition in positive ion mode. MS survey scans were acquired from 375-1500 m/z in profile mode at a resolution of 60,000. AGC target was set to 300%, allowing a maximum injection time of 25 ms. Peptides with charge states 2-6 were subjected to HCD fragmentation (fixed CE = 30%, AGC = 200%, maximum injection time 80 ms) and analysed in the Orbitrap at a resolution of 45,000. The isolation window was set to 1.4 m/z. Settings Lumos: Elution was carried out at a constant flow rate of 250 nL/min using a 180-min method: 8-33% solvent B (0.1% formic acid in 80% acetonitrile) within 120 min, 33-48% solvent B within 25 min, 48-98% buffer B within 1 min, followed by column washing and equilibration. Data acquisition was carried out using data-dependent acquisition in the positive ion mode. MS survey scans were acquired from 375-1500 m/z in the profile mode at a resolution of 120,000. AGC target was set to $4e^5$ charges, allowing a maximum injection time of 50 ms. Peptides with the charge states 2-5 were subjected to CID fragmentation (fixed CE = 35%, AGC = $2e^5$) and analysed in the Orbitrap at a resolution of 30,000. The isolation window was set to 1.2 m/z.

***Data-independent acquisition (DIA) proteomics of S. pyogenes ΔyebC strain***

10 µg of a 1:1 mixture of hydrophilic and hydrophobic carboxyl-coated paramagnetic beads (SeraMag, GE Healthcare) was added for each µg of protein. Protein binding was induced by the addition of acetonitrile to a final concentration of 70% (v/v). Samples were incubated for 10 min at room temperature. The tubes were placed on a magnetic rack, and beads were allowed to settle for three min. The supernatant was discarded, and beads were rinsed thrice with 80% ethanol. Beads were resuspended in a digestion buffer containing 50 mM triethylammonium bicarbonate (Sigma) and Trypsin and lys C (SERVA) in a 1:50 enzyme-to-protein ratio. Protein digestion was carried out for 14 h at 37°C. The peptide supernatant was then recovered and acidified with 2% ACN and 0.1% trifluoroacetic acid.

Label-free DIA analyses of peptides were acquired over 120 min by an Orbitrap Exploris 480 (Thermo Scientific) coupled to a 3000 RSLC nano UPLC (Thermo Scientific) from 1 µg of peptides. Samples were loaded on a pepmap trap cartridge (300 µm i.d. x 5 mm, C18, Thermo) with 2% acetonitrile, 0.1% TFA at a flow rate of 20 µL/min. Peptides were separated over a 25 cm analytical column (PepSep C18, 75 µm I.D., 1.5 µm). Solvent A consists of 0.1% formic acid in water. Elution was carried out at a constant flow rate of 250 nL/min within 120 min. Initially, a two-step linear gradient was applied: 3-30% solvent B (0.1% formic acid in 80% acetonitrile) within 70.5 min, 30-45% solvent B within 13 min, followed by column washing and equilibration. The column was kept at a constant temperature of 50°C.

The MS was operated in DIA mode for single-injection quantitative measurements of individual samples with the following settings: 60k MS1 resolution, MS1 scan range 375-1400 m/z, 15k MS2 resolution, MS2 scan range 120-1600 m/z, Normalized AGC target of 1000%, maximum injection time 54 ms, and fixed normalized collision energy of 30. 12 m/z precursor isolation windows with optimized window placements from 400.4319 to 1204.7975 m/z.

#### Preparation of NGS libraries

***RNA-seq libraries***

For each sample, 10 µg of RNA were treated with Turbo DNase (Thermo Scientific) and the RNA was purified using the RNA Clean & Concentrator-5 kit (Zymo Research) according to the “Total RNA Clean-up” protocol. Ribosomal RNA was depleted with Pan-Bacteria riboPOOL kit (siTOOLs) and RNA was once again purified using RNA Clean & Concentrator-5. 40 ng of RNA of each sample were used to create the libraries using the NEBNext Ultra II Directional RNA Library Prep Kit (New England Biolabs). The libraries were amplified with 10 cycles of PCR and twice purified with the NEBNext Sample Purification Beads to completely remove self-ligated adapters.

***iCLIP libraries***
The SMI RNA was dephosphorylated in 20 µL of MES-PNK buffer with 10 mM DTT and 15U of T4 PNK (Thermo Scientific) for 30 min at 37°C, purified with the Oligo Clean & Concentrator kit (Zymo Research) and eluted with 11 µL of mQ. L3-App oligo (Supplementary table 1) was ligated to all samples with T4 RNA Ligase 2, truncated K227Q (New England Biolabs) in the presence of 12% PEG8000 at 25°C for 2 h. RNA was resolved on 12% Urea TBE PAGE, the RNA fragments with the lengths 40-300 nt were cut and eluted from the gel. RNA was ethanol precipitated with GlycoBlue Coprecipitant and re-dissolved in 11 µL of mQ. Reverse transcription was performed with SuperScript III (Thermo Scientific) and RToligo primer and RNA was degraded with 0.2 M NaOH by incubation at 65°C for 15 min. cDNA was purified with Oligo Clean & Concentrator kit and eluted with 8 µL of mQ. The second adaptors L07-L12clip2.0 were ligated with T4 RNA Ligase 1, High Concentration (New England Biolabs) in the presence of 15% PEG8000 at 16°C overnight. For each sample 10 µL of MyONE Silane beads (Thermo Scientific) were washed with RLT buffer (Qiagen) and resuspended in 60 µL of RLT buffer. The 20 µL ligation reactions were mixed with 60 µL of the resuspended beads and 60 µL of ethanol and incubated for 10 min on the bench. The beads were washed three times with 900 µL of 80% ethanol and dried for 5 min on the bench. The purified cDNA was eluted from the beads with 18 µL of mQ, the beads were magnetized and the supernatants were collected. The cDNA libraries were amplified by a two-round PCR with Phusion DNA polymerase (Thermo Scientific). The first 10-cycle PCR was performed with primers P5Solexa_s and P3Solexa_s. The PCR products were resolved on 12% native TBE PAGE, the fragments 75-300 bp were excised and eluted from the gel. After ethanol precipitation, they served as the template for the second round of PCR with primers P5Solexa and P3Solexa. For the SMI control, 9 to 11 cycles of the PCR were performed, and for the other samples and controls, 16 cycles. The amplified libraries were purified using the ProNex Size-Selective Purification System (Promega) at a sample-to-ProNex (vol/vol) ratio of 1:2.2. Sequencing of the libraries was performed using the Seq_1 and Seq_2 primers in a paired-end mode.

***Ribosome profiling libraries***
The RNA was dephosphorylated with T4 Polynucleotide Kinase (Thermo Scientific) in buffer A for 2 h at 37°C. The RNA was precipitated with 2-propanol and GlycoBlue Coprecipitant and re-dissolved in 5.5 µL of mQ. The pre-adenylated adapter with unique molecular identifier (Supplementary table 1) was ligated with T4 RNA Ligase 2, truncated K227Q (New England Biolabs) in the presence of 25% PEG8000 at 25°C for 2 h. The adapter was also ligated to the mix of 15 and 45 nt control RNAs and this sample later served as guides for gel-purification. In order to remove the non-ligated adapter 0.5 µL of Rec J exonuclease (Biosearch Technologies) and 0.5 µL of yeast 5′-deadenylase (New England Biolabs) were added and the reactions were incubated at 30°C for 45 min. Reverse transcription was performed with SuperScript III (Thermo Scientific) and RT primer and the RNA was degraded with 0.2 M NaOH by incubation at 65°C for 15 min. cDNA was purified using the Oligo Clean & Concentrator kit and eluted with 8 µL of mQ. cDNA fragments were resolved on 10% Urea TBE PAGE. Using the size guides the fragments with 15 to 45 nt inserts were excised and isolated from the gel. cDNAs were precipitated with 2-propanol and GlycoBlue Coprecipitant and re-dissolved in 15 µL of mQ. Circularization of the cDNA molecules was performed with CircLigase II ssDNA Ligase (Biosearch Technologies) for 1 h at 60°C. The cDNA libraries were then amplified with primers introducing the sequencing barcodes. The amplified libraries were purified with the ProNex Size-Selective Purification System (Promega) at a sample-to-ProNex (vol/vol) ratio of 1:2.2.

#### Cloning, expression and purification of *E. coli* YebC proteins
The coding sequences of *E. coli yebC* paralogs *b1983* and *b1864* were amplified from genomic DNA of strain BW25113 with primers listed in Supplementary table 1. The expression vector pET-21a(+) was amplified with primers excluding the T7 tag sequence. HiFi assembly (New England Biolabs) was

used to assemble the expression vectors. The HA tag was then introduced to the C-termini of the genes by the whole-plasmid PCRs and circularization with the KLD mix (New England Biolabs). The mutations M2 and Y84A were introduced to the b1983 sequence by the whole-plasmid PCRs and circularization with KLD mix. The introduction of mut-2 required two cycles of mutagenesis.

The plasmids pET-21a_b1983-HA, pET-21a_b1983-mut2-HA, pET-21a_b1983-Y84A-HA and pET-21a_b1864-HA (Supplementary table 2) were introduced by transformation into *E. coli* BL21(DE3) cells (Thermo Scientific). The colonies were inoculated into 200 mL of LB medium and the cells were grown at 37°C with 180 rpm shaking until $OD_{600\ nm}$ = 0.6. The cultures were cooled on ice, the protein expression was induced with 200 µM IPTG overnight at 25°C with shaking at 180 rpm. The cells were collected by centrifugation and stored at -80°C. The cells were lysed with the B-PER Complete Bacterial Protein Extraction Reagent (Thermo Scientific) supplemented with 500 mM NaCl, 20 mM imidazole and cOmplete, EDTA-free Protease Inhibitor Cocktail (Merck). The proteins were immobilized on HisTrap HP columns (GE Healthcare), washed with buffer A (50 mM Tris-HCl pH 7.4, 500 mM NaCl, 20 mM Imidazole) and eluted with buffer B (50 mM Tris-HCl pH 7.4, 500 mM NaCl, 500 mM Imidazole) using a linear gradient on the ÄKTA pure chromatography system (Cytiva). The proteins were further purified by gel-filtration on Superdex 200 13/300 increase columns (Cytiva) in SEC buffer (25 mM Tris-HCl pH 7.4, 200 mM NaCl, 0.5 mM EDTA, 2 mM DTT, 10% glycerol). The purified proteins were analyzed by Coomassie-stained SDS-PAGE, aliquoted and stored at -80°C.
